# Individual Differences in Explicit Strategy Development During Visuomotor Adaptation: The Influence of Perturbation Magnitude

**DOI:** 10.64898/2026.04.14.718592

**Authors:** Elysa Eliopulos, Denise Y.P. Henriques, Bernard M. ’t Hart

## Abstract

**Background:** Humans rapidly adapt their movements to compensate for changes in the environment, yet how explicit movement strategies emerge during adaptation remains poorly understood. These strategies have traditionally been described by using a smooth, exponential learning curve, suggesting a relatively uniform pattern of strategy development across individuals (Taylor et al., 2014; McDougle et al., 2015). However, recent results suggest substantial variability in how individuals develop explicit aiming strategies, such as abrupt transitions (D’Amario et al., 2024; Townsend et al., 2026), or exploratory periods, or delayed stabilization of compensatory behaviour (Tsay et al., 2024), and some participants never developing a strategy. These findings raise the possibility that smooth group-averaged learning curves may obscure meaningful heterogeneity in individual strategy development. This study therefore examines whether explicit strategy development follows distinct individual-level patterns and whether these dynamics are influenced by perturbation size.

**Methods:** Participants completed a visuomotor rotation reaching task where a cursor they controlled with their unseen, right hand was rotated around the home position using one of five perturbations (20°, 30°, 40°, 50°, and 60°). Before each reach, participants reported their explicit strategy by aiming an arrow to indicate their planned hand movement direction.

**Results:** Based on previous findings, we hypothesized that participants would exhibit at least three distinct patterns of explicit strategy development emerged: 1) gradual/incremental, 2) stepwise, and 3) exploratory. Instead, participants were distributed along a continuum of strategy development profiles, with different regions along this continuum resembling these previous proposed patterns. Where learners fell along this continuum varied with perturbation magnitude, such that larger rotations associated with greater exploratory behaviour and proportion of strategy-use.

**Conclusion:** Our findings reveal substantial heterogeneity in how explicit strategies emerge during visuomotor adaptation and suggest that smooth group-averaged learning curves may obscure important individual differences in learning dynamics. Notably, perturbation magnitude appears to systematically influence where individuals fall along this continuum, providing a potential mechanism for understanding how explicit strategies emerge and stabilize over time.

## Introduction

Humans continuously adapt their movements in response to changes in the body and environment. This ability, known as motor adaptation, allows movements to remain accurate and coordinated despite altered sensory feedback or changing task demands. Examples from daily life include golfing on tilted surfaces or walking on an icy sidewalk, where altered object dynamics initially produce discrepancies between intended and actual movement outcomes.

Individuals may deliberately aim their putt away from the hole or take shorter steps to compensate for these discrepancies. With practice, the resulting movement errors are gradually reduced, resulting in smoother and more accurate movements.

A common method for studying motor adaptation is the visuomotor rotation (VMR) paradigm, where visual feedback of the hand is rotated relative to the intended movement direction (Fig. 1). During adaptation, the visual feedback of the unseen hand, (path is indicated as dashed lines in Fig 1), i.e. cursor, is rotated relative to the actual hand movement direction (solid line, in Fig 1, ii). As a result, participants initially experience systematic reaching errors, but with training they gradually adjust their movements to compensate for the perturbation (iii). Following removal of the rotation, reach aftereffects are often observed, indicating that adaptation-related changes persist beyond the perturbation itself (iv).

**Figure 1.**
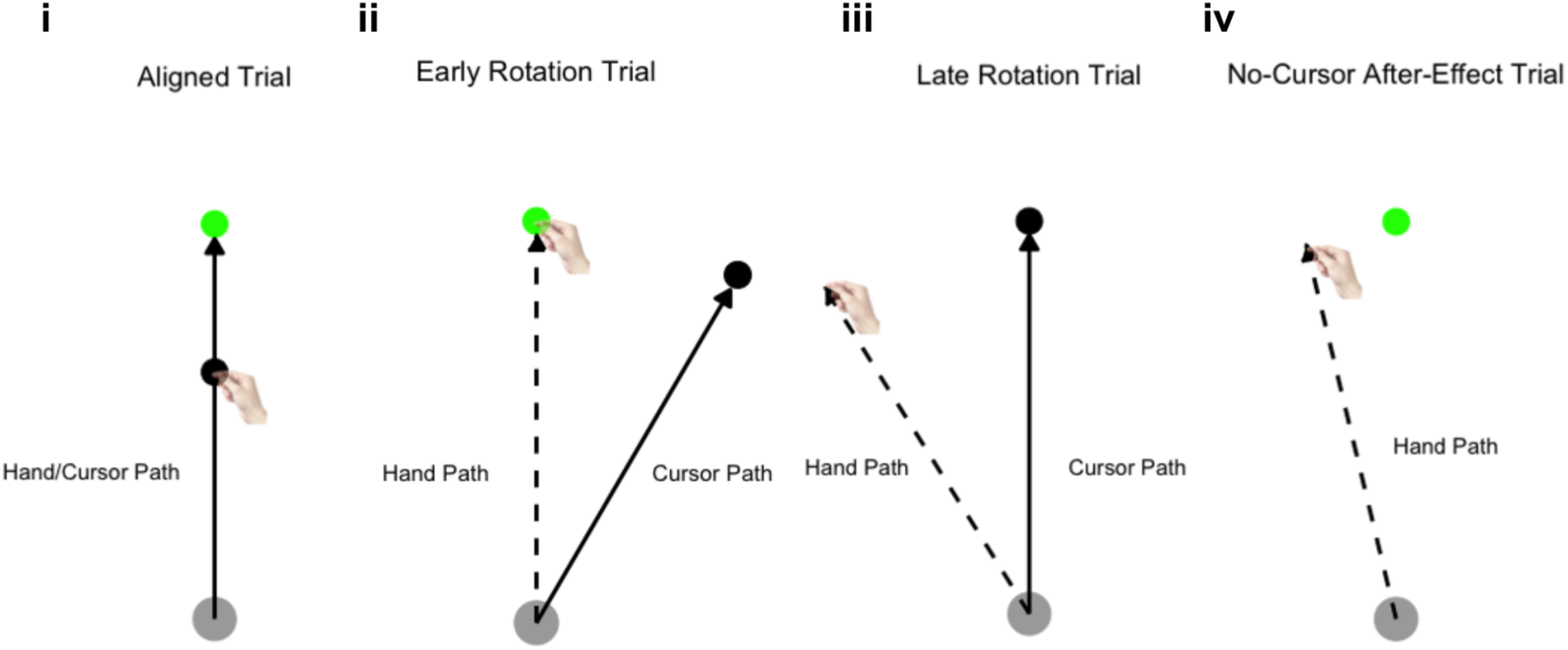
Visuomotor Adaptation Paradigm. Participants are tasked with reaching a cursor straight to a green target (aligned trial). At some point in time, a cursor rotates from the participant’s initial hand path (early rotation trial). A participant can counter this cursor rotation by reaching their hand in a direction that is opposite from the perturbation path (late rotation trial approximately after 20 trials). As participants adapt to the rotation, this modified behaviour may still be apparent following removal of the cursor rotation (no-cursor after-effect trial).

Explicit adaptation refers to consciously applied movement strategies that individuals use to reduce movement errors and compensate for perturbations. Explicit adaptation has traditionally been characterized using smooth exponential learning curves averaged across participants (Taylor et al., 2014; McDougle et al., 2015; Brudner et al., 2016). However, averaging behaviour across individuals may obscure important variability in how these strategies develop. Recent studies suggest that explicit strategy development can involve abrupt transitions (D’Amario et al., 2024; Townsend et al, 2026) or exploratory periods, or delayed stabilization of compensatory behaviour (Tsay et al., 2024). Together, these findings raise the possibility that smooth group-averaged learning curves may not accurately capture the dynamics of explicit strategy development at the individual level. In the current study, we characterize how explicit strategies emerge during visuomotor adaptation and examine whether their development varies as a function of perturbation magnitude.

## Explicit and Implicit Adaptation

Behavioral changes during visuomotor changes are thought to reflect contributions from both implicit and explicit processes (Heuer & Hegele, 2007; Sülzenbrück & Heuer, 2009; Benson et al., 2011; Taylor et al., 2014; Bond & Taylor, 2015; Neville & Cressman, 2018). Implicit adaptation is generally considered an automatic and unconscious recalibration process that can be observed through aftereffects during no-cursor trials following removal of the perturbation (fig 1, iv). In contrast, explicit adaptation involves consciously applied movement strategies that can be voluntarily adjusted to counter the rotation. Overall adaptation performance, typically measured as changes in the direction of hand movements across training, reflects contributions from both processes (Fig. 2). Within this framework, explicit strategies are thought to contribute strongly to rapid early error reduction, whereas implicit recalibration develops more gradually over time (Taylor et al., 2014; McDougle et al., 2015). Although both processes contribute to visuomotor adaptation, recent evidence suggests that their relationship may not be linearly additive (’t Hart et al., 2024). Understanding how these processes interact remains an active area of research and highlights the importance of characterizing the development of explicit strategies during adaptation.

**Figure 2.**
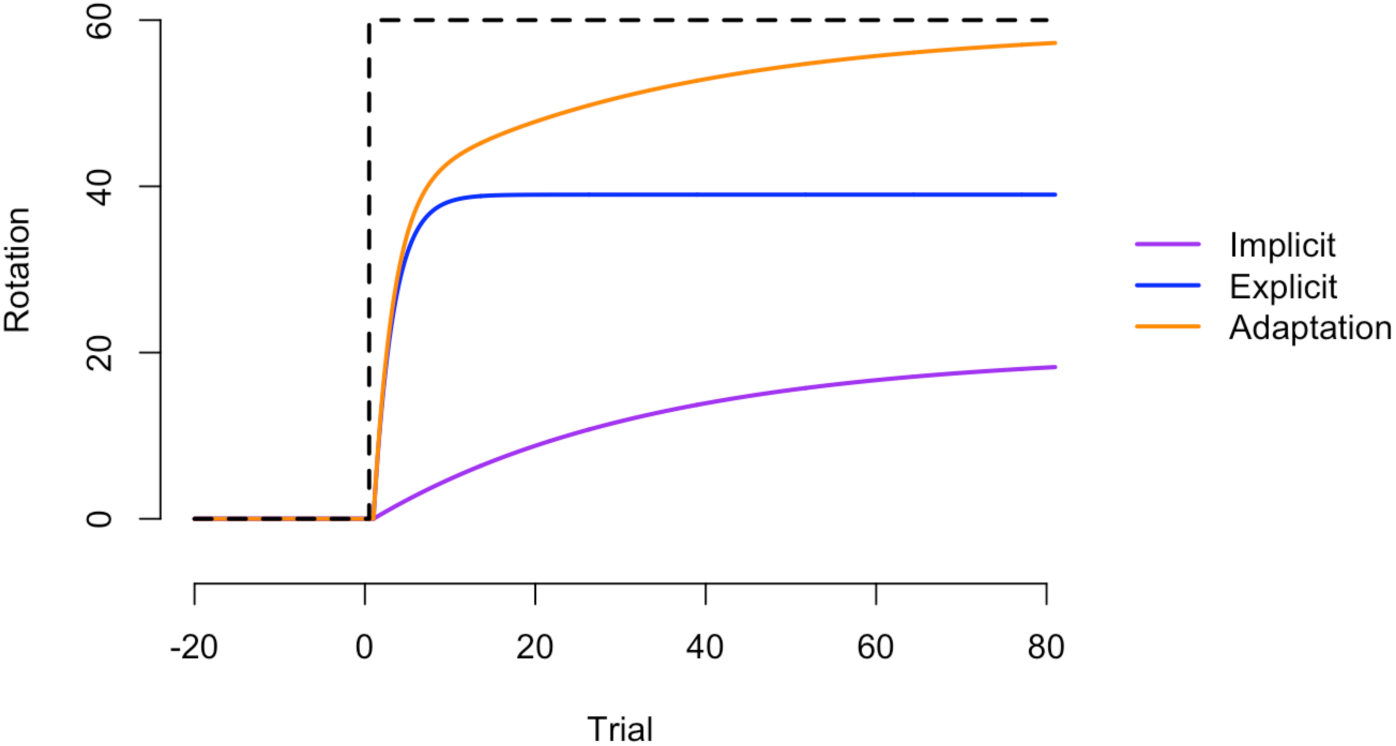
Model of Visuomotor Adaptation Timecourses. Total adaptation (orange) reflects overall changes in reach behaviour during adaptation. Explicit adaptation (blue) and implicit adaptation (purple) represent the canonical timecourses of these two subcategories of motor adaptation.

## Measuring Explicit Adaptation

Explicit adaptation reflects consciously applied movement strategies, and several methods have been developed to quantify intended aiming direction during visuomotor rotation adaptation. One widely used approach involves aiming reports, in which participants indicate their intended reach direction prior to movement (Hegele & Heuer, 2010; Taylor et al., 2014). For example, Taylor et al. (2014) uses a circular ring of numbered landmarks from which participants selected their intended aiming direction as shown in Figure 3a (McDougle et al., 2016). More recently, continuous aiming-report paradigms have been developed in which participants indicate their intended movement direction by rotating an on-screen arrow or marker prior to each reach (Bindra et al., 2021; Hadjiosif et al., 2024; D’Amario et al., 2024).

**Figure 3.**
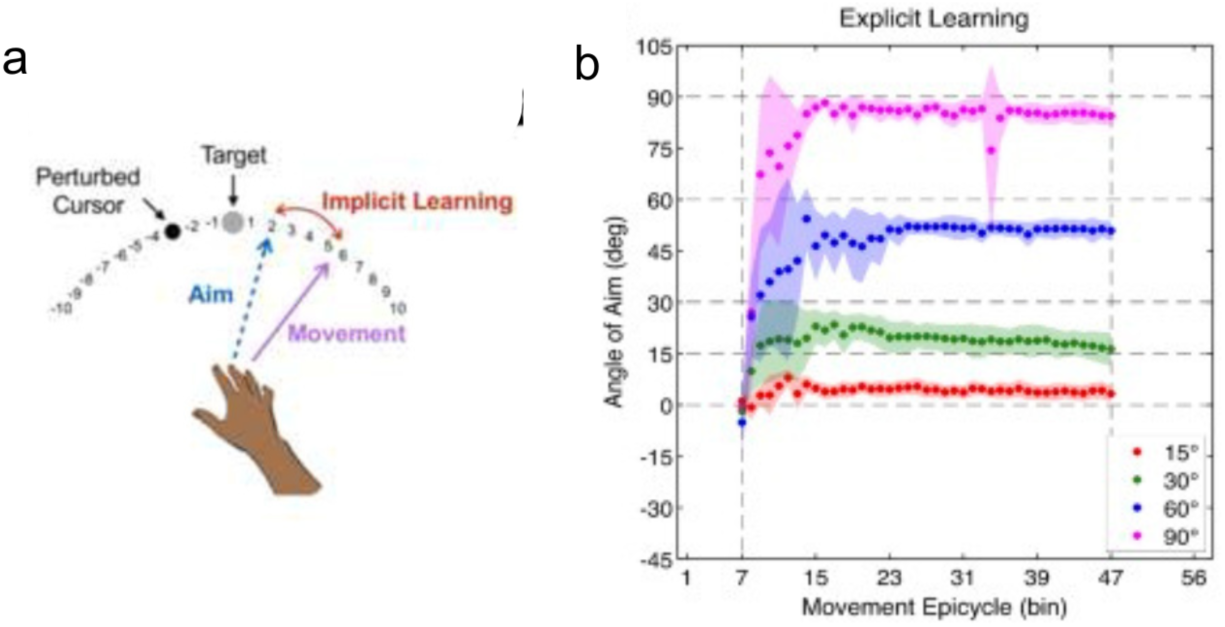
Explicit Visuomotor Adaptation. This process is assessed via re-aiming reports. **A:** Derived from McDougle et al (2016). The verbal report method for measuring explicit strategies to counter a rotation. Participants point to a numbered landmark that indicates the direction they are moving their hand in. **B:** Derived from Bond & Taylor (2015), the degree of aim is dependent on rotation size. When a participant is introduced to a 60-degree counterclockwise rotation, they will closely adjust their aim 60-degrees clockwise.

One other approach infers explicit adaptation indirectly by instructing participants to implement or suppress learned strategies (Werner et al., 2015; Neville & Cressman, 2018; Modchalingam et al., 2019; Gastrock et al., 2020; Heirani Moghaddam et al., 2021). While this method uses reaches to report strategy and may hence be closer to what participants do during the adaptation trials, this requires subtraction, which may not work (’t Hart et al., 2024) and multiple trials for a single measurement, such that it cannot easily provide a continuous measure of explicit strategy development. Another approach is to use delayed feedback to eliminate implicit contributions and simply look at the adaptation timecourse as a direct measure of explicit adaptation (Schween & Hegele, 2017; Tsay et al., 2023; Ding et al., 2026). However, implicit aftereffects can still be observed in many studies using delayed feedback (Brudner et al., 2016; Avraham et al., 2019; McDougle & Taylor, 2019; Tsay et al., 2023). This implies that delayed feedback does not result in a pure measure of explicit adaptation. This is why we opt here to use interleaved aiming as a reliable and direct way to obtain a continuous measure of explicit strategy throughout adaptation.

Final strategy magnitude scales with perturbation size, with larger rotations typically eliciting greater explicit compensation (Bond & Taylor, 2015; Tsay et al., 2024; Figure 3b). This relationship raises the possibility that perturbation magnitude may influence not only the amount of explicit compensation expressed by participants, but also the manner in which explicit strategies develop over training. Consequently, trial-by-trial aiming reports make it possible to determine whether perturbation magnitude influences not only the amount of explicit strategy expressed, but also the timecourse by which explicit strategies emerge during adaptation.

## Individual Differences in Explicit Strategy Development

Although explicit strategy development has traditionally been characterized using smooth exponential learning curves (Taylor et al., 2014; McDougle et al., 2015; Brudner et al., 2016), recent work has begun investigating the underlying structure of explicit strategy development.

D’Amario et al. (2024; Experiment 4) examined trial-by-trial aiming reports and observed considerable divergence from the smooth exponential timecourses typically associated with explicit strategy development. Most participants exhibited abrupt transitions into stable aiming solutions, whereas others showed little evidence of explicit aiming despite successfully adapting their reaches to the perturbation (Fig 4). Interestingly, despite these markedly different individual trajectories, the group-averaged time courses for total, implicit, and explicit adaptation all resembled smooth exponential learning curves. While averaging appropriately captures the gradual nature of total and implicit adaptation, it may obscure the nature of explicit strategy development as well as meaningful differences between strategy development of individual participants.

**Figure 4.**
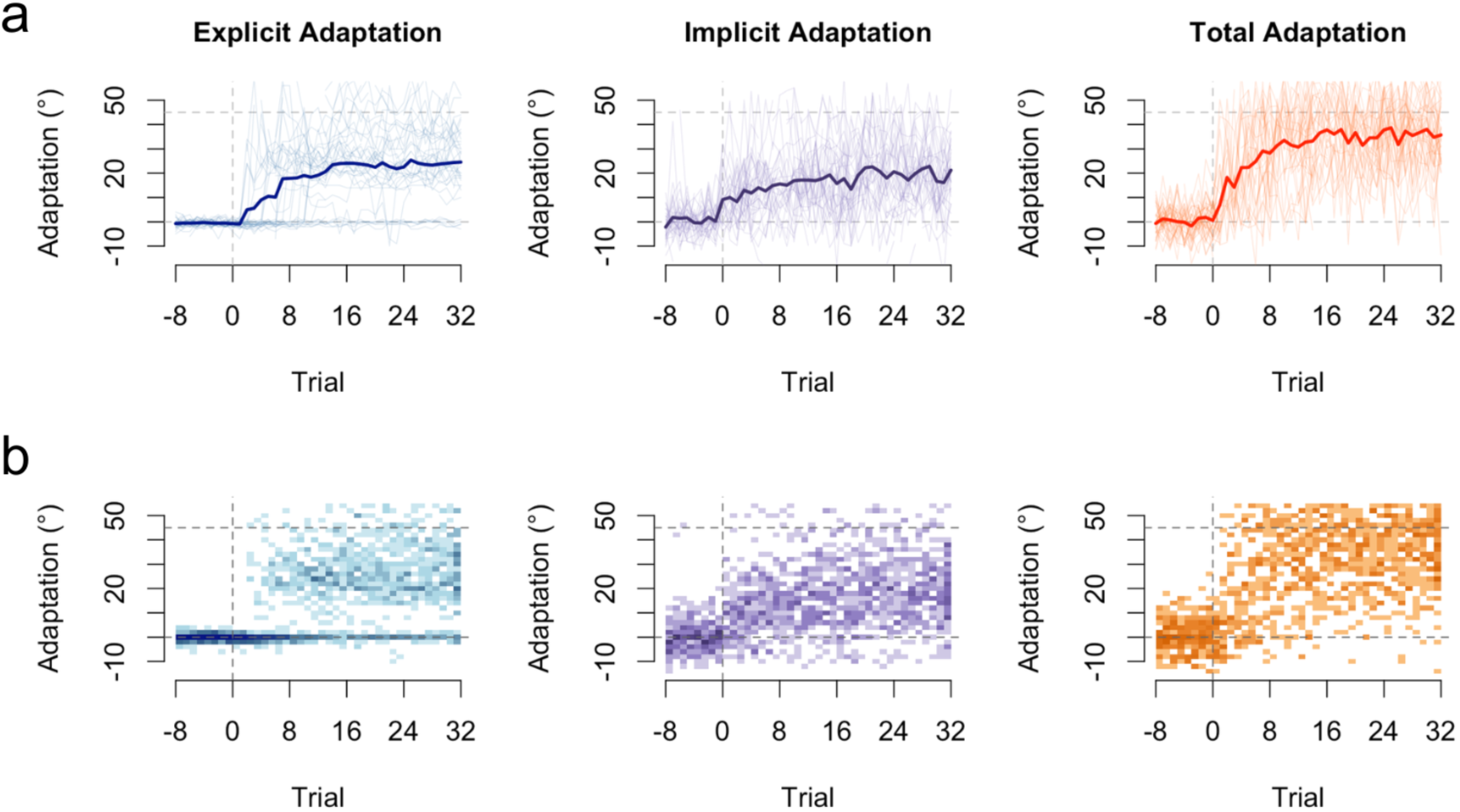
Overall, Implicit, and Explicit Adaptation Timecourses. Figure adapted from D’Amario et al., 2024. **A:** Individual reach deviations. Thin lines represent individual participants, darker lines represent group means for reaches. Left panel displays explicit adaptation, measured with arrow aiming. Middle panel shows implicit reaches, measured with no-cursor reach trials, Right panel reflects total adaptation. **B:** 2d histograms of explicit (left panel), implicit (middle panel) and total (right panel) adaptation. Darker regions reflect areas of common responses. Note that the timecourses for overall and implicit adaptation, while noisy, look more homogenous, while for explicit adaptation, most participants start indicating a strategy rather suddenly, leaving a gap in the data at around 10 degrees, and that some participants never indicate a strategy, causing a small region of aiming to remain at around 0 degrees. Since these two properties are not present in total a group averaging may not be capturing the underlying structure of the explicit process.

Similar abrupt transitions in strategy development have been observed in a simplified web-based aiming task, where participants adjusted the direction of a cannon using button presses to hit a target (Townsend et al., 2025), with most participants exhibiting abrupt shifts into stable aiming solutions. Likewise, Tsay et al. (2024) highlighted substantial variability in explicit aiming behaviour across studies, including abrupt shifts in strategy use, slower strategic adjustments, and exploratory responding. Additional evidence for individual differences comes from a study examining how participants generalize learned strategies. Ding et al. (2026) used cluster-based analyses in a web-based visuomotor adaptation task and identified variability in the compensatory rules participants applied to probe targets following adaptation. The authors interpreted these findings as evidence that individuals may adopt different strategic approaches when solving a perturbation. However, because these approaches infer strategy indirectly at non-adapted targets, it remains unclear whether similar patterns would be observed using continuous measures of explicit aiming at the trained targets.

Collectively, these findings (D’Amario et al., 2024; Townsend et al., 2025; Tsay et al., 2024; Ding et al., 2026) suggest substantial heterogeneity in explicit strategy development. That is, to understand the process of explicit strategy development, we should take individual, continuous aiming report timecourses into account. However, it remains unclear whether the observed patterns reflect discrete learning “phenotypes” or regions along a broader continuum of strategy development. Furthermore, little is known about the factors that influence where individuals fall within this space. One potential factor is perturbation magnitude, which may affect both the likelihood of strategy use and the manner in which explicit strategies emerge over the course of learning.

## The effect of perturbation size on strategic and implicit learning

The substantial variability in explicit strategy development described above raises the question of what factors contribute to these individual differences. One candidate factor is perturbation magnitude. Previous work has shown that the magnitude of explicit compensation generally scales with rotation size, such that larger rotations elicit larger aiming adjustments (Bond & Taylor, 2015, see Fig 3B). In addition, recent findings suggest that larger perturbations magnitude may influence whether participants develop an explicit strategy at all. For example, D’Amario et al., (2024) observed that participants exposed to smaller rotations often exhibited little measurable aiming behaviour. Participants exposed to larger rotations were more likely to elicit more bi-modal patterns of explicit strategy use: some participants apparently did not have a strategy, while others did. These findings raise the possibility that perturbation magnitude influences not only the amount of explicit compensation expressed by participants, but also if and how explicit strategies emerge over the course of learning. Larger perturbations may increase the likelihood of exploratory behaviour, abrupt strategy discovery, or explicit strategy use, whereas smaller perturbations may be more likely to produce little to no measurable aiming compensation.

In contrast, implicit adaptation appears considerably less sensitive to perturbation magnitude. Across a wide range of rotation sizes, implicit aftereffects typically remain constrained to approximately 10–20° despite substantial increases in perturbation size (Bond & Taylor, 2015; Morehead et al., 2017; Kim et al., 2018; Modchalingam et al., 2019, but see: Modchalingam et al., 2023). Consequently, whereas explicit compensation tends to scale with perturbation magnitude, implicit adaptation appears to reach a relatively fixed asymptote. This dissociation suggests that perturbation size may primarily influence strategic aspects of adaptation.

Although perturbation magnitude influences average levels of explicit compensation, it remains unclear whether it also influences the development of explicit strategies at the individual level. For example, larger perturbations may increase the likelihood of exploratory behaviour, abrupt strategy discovery, or explicit strategy use, whereas smaller perturbations may promote more incremental adjustments or little measurable aiming. The present study therefore examines how perturbation magnitude influences individual differences in explicit strategy development and whether these differences are associated with subsequent implicit aftereffects.

## Insight Learning as a Potential Mechanism for Explicit Strategy Development

One possible explanation for abrupt changes in explicit aiming behaviour is insight learning. Rather than developing incrementally, explicit strategies may sometimes emerge suddenly following the discovery of a successful solution to the perturbation. Such abrupt improvements are often described as “Aha!” moments and have been observed across a variety of problem-solving tasks (Metcalfe & Wiebe, 1987; Danek et al., 2020).

Dubey and colleagues proposed that insight arises when individuals solve a problem sooner than expected, generating a metacognitive prediction error and a corresponding feeling of sudden solvability. Applied to visuomotor adaptation, participants may initially struggle to determine how to compensate for a perturbation before suddenly discovering an aiming strategy that substantially reduces movement error. Such a discovery would be expected to produce an abrupt change in explicit aiming behaviour rather than an incremental increase in compensation.

Consistent with this possibility, recent studies using continuous aiming reports have documented sudden transitions in explicit strategy use during visuomotor adaptation (D’Amario et al., 2024; Townsend et al., 2026). These abrupt shifts are consistent with the notion that some participants may discover a compensatory solution suddenly rather than gradually refining an existing strategy. However, whether such transitions represent a distinct form of learning or one region along a broader continuum of strategy development remains unclear.

## Proposed Types of Explicit Learning Timecourses

Recent studies suggest that explicit strategies may emerge through a variety of learning trajectories rather than a single smooth timecourse. Here, we consider three candidate patterns of explicit strategy development that capture distinct hypotheses about how explicit strategies may emerge during adaptation.

One possibility is an incremental strategy-development pattern, in which aiming adjustments increase more or less monotonically across trials. This type of timecourse would be most consistent with the smooth exponential learning curves traditionally used to characterize explicit strategy development.

A second candidate pattern is a stepwise transition of strategy development. In this case, participants may exhibit a relatively abrupt transition from little or no explicit compensation to a stable aiming solution. Such abrupt changes are consistent with observations from recent aiming-report studies (D’Amario et al., 2024; Townsend et al., 2026) where they have been interpreted as reflecting the sudden discovery of a successful compensatory strategy.

A third possibility is an exploratory pattern, in which participants test multiple potential solutions before settling on a stable aiming strategy. This behaviour would be characterized by elevated trial-to-trial variability and fluctuations in aiming direction prior to stabilization.

Exploratory behaviour has been highlighted as a potential feature of explicit adaptation in recent reviews of aiming-based measures (Tsay et al., 2024). Figure 5 illustrates the visual patterns of these proposed strategy development types.

**Figure 5.**
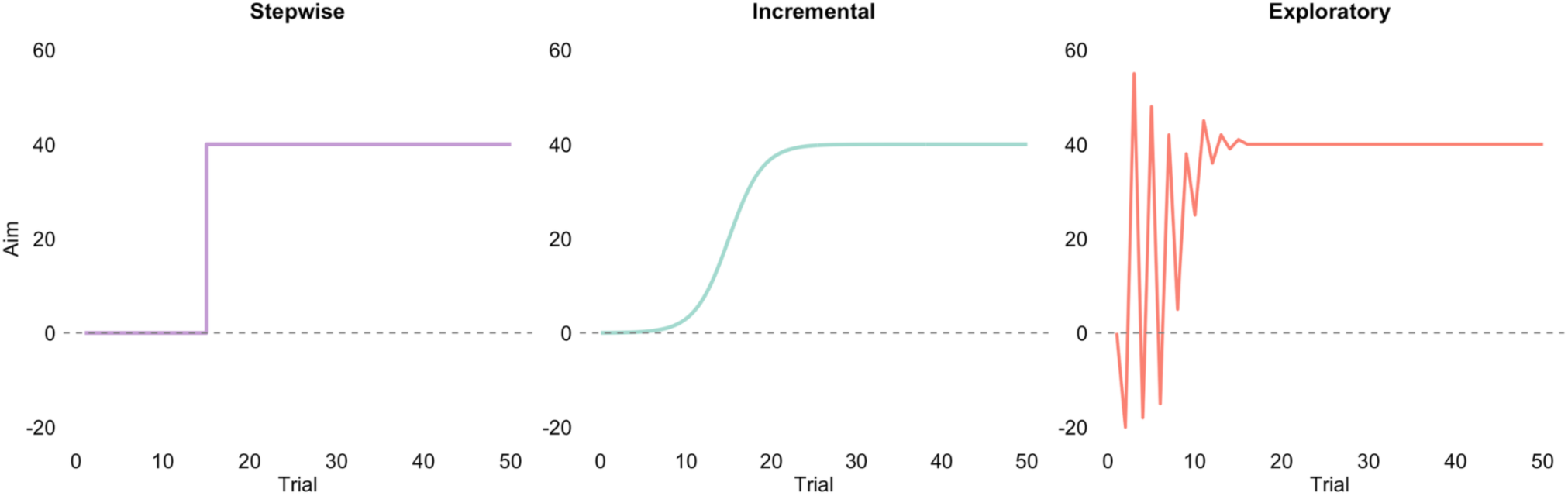
Model of Proposed Strategy Development Types. Left panel illustrates an Aha! Moment, characterized by an abrupt shift in aiming, based on work by D’Amario et al. (2024) and Townsend et al. (2025). Middle panel models a sigmoidal pattern as one example of a slower. incremental strategy development pattern. Right panel illustrates exploratory patterns, characterized by variable trial-to-trial adjustments, based on work by Tsay et al. (2024).

Other patterns may be possible as well. Importantly, it remains unclear whether these proposed patterns, or any others, represent distinct learning phenotypes or are simply points along a broader continuum of strategy development patterns. Consequently, the present study examines individual aiming timecourses without assuming a priori that participants belong to discrete categories.

## Objectives and Hypothesis

The current project aims to characterize individual differences in in explicit strategy development during visuomotor adaptation using continuous trial-by-trial aiming reports. In particular, we sought to determine whether explicit strategy development follows a common learning trajectory across individuals or whether substantial heterogeneity exists in how participants discover and implement compensatory strategies.

A second objective is to determine whether perturbation magnitude influences the development of explicit strategy. Because larger rotations typically elicit great compensation, and larger rotation are more salient, we expect that perturbation magnitude would influence both the likelihood of strategy use and the way the strategies are developed. Specifically, larger perturbations are expected to increase the prevalence of strategy use and promote more abrupt or exploratory forms of strategy development. We may particularly see an increased likelihood of “Aha!” learning with increasingly salient errors.

Finally, we examined whether individual differences in explicit strategy development were associated with differences in subsequent implicit aftereffects. Given previous evidence that explicit and implicit adaptation interact, we explored whether variation in strategy development would be related to differences in the magnitude of implicit adaptation.

Based on previous findings, we expected individual aiming trajectories to exhibit substantial heterogeneity. We further anticipated that some trajectories would resemble incremental, stepwise, or exploratory patterns of strategy development. However, because it remains unclear whether such patterns reflect discrete learning phenotypes or regions along a broader continuum, no specific predictions were made regarding the number or nature of resulting strategy groups.

## Methods

### Participants

Two hundred and twelve right-handed undergraduate students from York University were recruited through the Kinesiology Undergraduate Research Experience (KURE) and Undergraduate Research Participant Pool (URPP) systems and received course credit for participation. Prior to participation, all individuals provided informed consent and completed a demographic questionnaire.

Participants were assigned to one of five visuomotor rotation groups (20°, 30°, 40°, 50°, or 60°). The study was designed to obtain 40 learners in each rotation group. A learner was defined a priori as a participant who compensated for at least 50% of the imposed rotation during the final 16 perturbation trials. Participants who did not meet this criterion were classified as non-learners and were not included in subsequent analyses. Recruitment continued until 40 learners had been obtained in each rotation group, resulting in a final sample of 200 learners (40 per group) from the original sample of 212 participants. All procedures complied with institutional and international ethical guidelines and were approved by York University’s Human Participants Review Committee.

## Apparatus

Participants sat in front of a digitizing tablet (Wacom Intuos Pro Large) while holding a stylus in their dominant hand. The hand workspace was occluded from direct view by a horizontal mirror positioned 26 cm above the tablet and 26 cm below a computer monitor (1680 × 1050 pixels; 60 Hz) (Figure 6A). Visual stimuli were projected onto the monitor and viewed as mirror reflections, creating the perception that stimuli were located in the same plane as the tablet surface.

**Figure 6.**
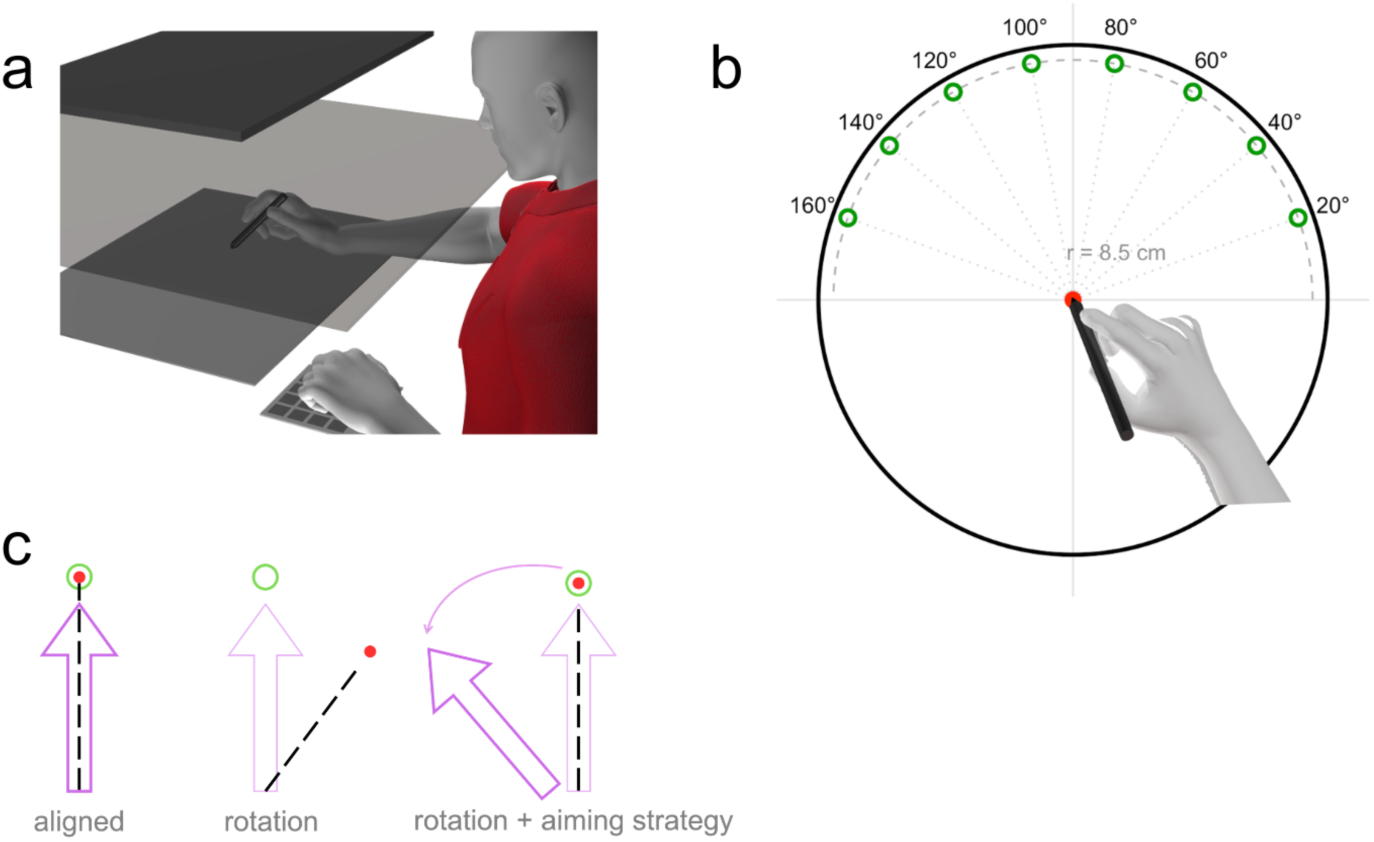
Experiment set up and task features. **A:** Participants sat in front of a digital tablet, holding a stylus on a tablet with their visually occluded hand. A mirror is placed over the tablet where participants can see the reflection from the monitor that is placed above the mirror. **B:** Target Locations. 8 Targets are placed at various locations at 20°, 40°, 60°, 80°, 100°, 120°, 140°, and 160°. Only one of 8 targets is presented at each reaching trial. **C:** Aiming Method. One of the 8 targets illustrated in Figure 6b is shown to the participant. They must change direction of the purple arrow by using the left and right arrow keys to indicate where they aim to move their hand to get the cursor straight to the target. The direction they point the arrow in is taken as an estimate of their conscious strategy to counter the visuomotor rotation.

Eight target locations were positioned at 8 cm distance and 20°, 40°, 60°, 80°, 100°, 120°, 140°, and 160° relative to the start position. One of each target was presented in random order across trials (Figure 6B). An 8.5-cm radius circular stencil was placed on the tablet to guide movement extent and constrain reaches to just beyond 8 cm.

## Task Procedure

Participants performed center-out reaching movements from a central home position toward one of eight target locations (Fig. 6B) using a digital stylus held in their right hand. Hand position was represented by a red cursor (0.5 cm diameter) displayed on the monitor. Each trial began when participants aligned the cursor with a white home position and maintained this position for 333 ms, after which the home position disappeared and a green target appeared, signaling movement initiation.

Before each reach, participants reported their intended movement direction using an aiming-report procedure (Fig. 6C). Specifically, participants rotated a purple on-screen arrow using left and right arrow keys operated with their left hand to indicate the direction they intended to move their hand. To improve sensitivity to explicit strategy use, the arrow was initially positioned 10° clockwise from the target. Participants without an explicit strategy would therefore rotate the arrow toward the target, whereas participants using a compensatory strategy would rotate the arrow beyond the target in the direction opposite the (clockwise) perturbation.

Following the aiming report, participants completed an 8-cm outward reaching movement. During cursor-feedback trials, the reach was completed as soon as the center of the cursor gets within 0.25 cm of the target’s center. After target acquisition, participants returned the stylus to the home position. During the return movement, a blue circle centered on the home position provided distance feedback, shrinking as the stylus approached the start location. The red cursor reappeared within 1.2 cm from the home position to facilitate accurate repositioning before the next trial.

## Experimental Schedule

The experimental schedule is illustrated in Figure 7. The task consisted of aligned cursor-feedback trials (light blue blocks), no-cursor trials (dark blue blocks), error-clamp trials (grey blocks), and a visuomotor rotation phase (pink block). Participants first completed 24 aligned trials (light blue block), followed by 16 no-cursor trials (dark blue block) to familiarize participants with the task. They then completed eight error-clamp trials with large cursor offsets (±80° and ±100°) to reinforce the distinction between hand and cursor movement and ensure participants understood the aiming-report procedure. After these trials, participants completed additional aligned and no-cursor blocks (8 trials each), which provided the baseline measures used to quantify changes during the subsequent adaptation phases. Prior to the final aligned block, participants received a reminder regarding the purpose of the aiming-report procedure (“Aim Reminder” in Fig. 7). Following this reminder, participants completed a final 8-trial aligned block before entering the rotation phase.

**Figure 7.**
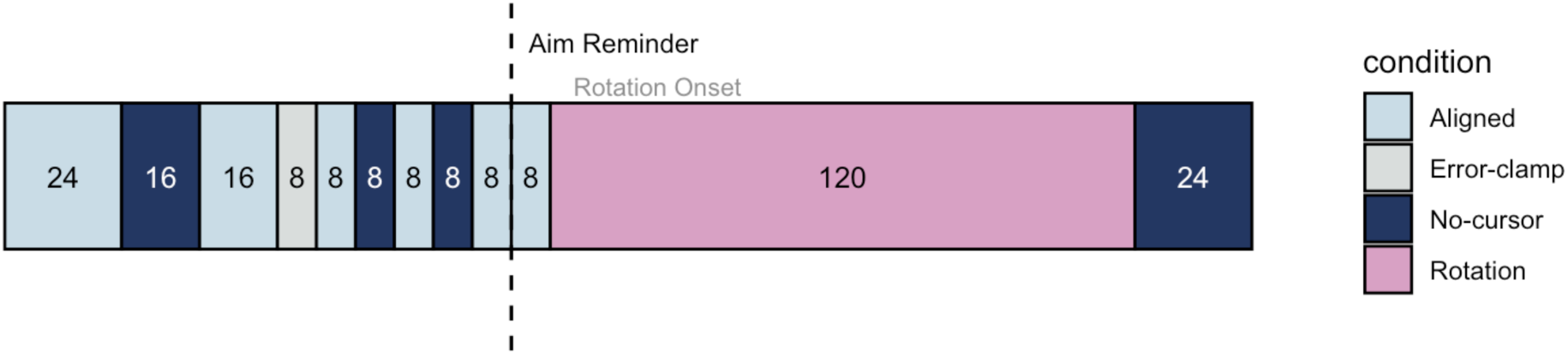
Experimental Task Schedule. Participants completed aligned, no-cursor, and error- clamp trials during the aligned phase. Prior to the rotation phase, participants received a final reminder regarding the aiming-report procedure. During the rotation phase, participants adapted to one of five visuomotor rotations (20°, 30°, 40°, 50°, or 60°) while reporting their intended aiming direction before each reach. The final no-cursor block was used to assess implicit aftereffects. Numbers indicate the number of trials in each block.

During the rotation phase (pink block in Fig. 7), participants completed 120 trials under one of five visuomotor rotations (20°, 30°, 40°, 50°, or 60°). Participants reported their intended aiming direction before every trial and were instructed to use whatever strategy was necessary to move the cursor to the target. Following adaptation, participants completed a final 24-trial no- cursor block (dark blue block) to assess implicit aftereffects.

## Instructions

Before beginning the experimental task, participants were informed that the cursor might not always follow their hand movement. During error-clamp trials (grey blocks in Fig. 7), participants were reminded that “the cursor is not your hand” and were instructed to move their hand slowly and ignore the cursor to observe the distinction between hand movement and cursor feedback. The goal of this dissociation is to set participants up for strategy reporting.

During cursor-feedback trials (light blue and pink blocks in Fig. 7), participants were instructed to use any strategy necessary to move the cursor to the target. During no-cursor trials (dark blue blocks in Fig. 7), participants were instructed not to use a strategy and instead move their hand directly to the target. Prior to the rotation phase, participants were reminded of the purpose of the aiming arrow (Fig. 6C) and were asked to explain, in their own words, what the arrow represented to ensure understanding of the task. These extensive instructions were provided before the rotation phase, such that no additional instructions should be necessary during the rotation phase. This should minimize experimenter influence on the natural development of explicit strategies.

## Analysis

All analyses were conducted using trial-by-trial aim deviation and reach deviation relative to target direction. Aim deviation, defined as the angular orientation of the aiming arrow relative to the target, served as a direct measure of explicit strategy use. Reach deviation, measured from the first sampled hand position occurring after the unseen hand exceeded a radial distance of 2 cm from the home position. The angular deviation of this position relative to the target direction served as an estimate of overall adaptation during trials with visual feedback and of implicit adaptation during no-cursor trials. Reach deviation during visual-feedback trials were baseline-corrected using the corresponding aligned baseline block immediately preceding perturbation onset (visual-feedback trials) or the final aligned no-cursor block (no-cursor trials). A one-way ANOVA was used to verify that overall adaptation during the late rotation phase increased with perturbation magnitude.

Only participants classified as learners (N=200, see Participants) were included in the data analyses. To determine whether learners developed an explicit aiming strategy, a confidence interval approach was applied to aiming behaviour during the final 16 perturbation trials.

Because baseline reaching variability is usually approximately 5° (Henriques et al., 2026), participants were required to have a lower bound of the 95% confidence interval larger than 5° to be considered as having a strategy. Participants without an explicit strategy would be expected to aim near the target location (∼0°), whereas participants using a compensatory strategy would systematically deviate their aim in the direction opposite the perturbation. We excluded one strategy-user from analysis as their aiming pattern never stabilized in the compensatory direction throughout the rotation phase. We next set out to understand the between-participant heterogeneity of aiming timecourses. The first step was to isolate participant strategy learning phases. Visual expertise was used to label onset and stabilization/end trials during strategic aiming. Strategy onset was visually identified as the first increase approximately around ±5°.

This 5° value ensures that arrow deviation is visually salient enough to appear off-target, reflecting conscious awareness of modified hand trajectories. The timepoint of strategy stabilization was identified as the period when trial-by-trial aiming variability reached a stable minimum, indicating convergence toward a common aim location.

The labeled strategic learning phases were next used as input to extract features of the strategy-development phase. These features were determined a priori with the expectation that distinct clusters as described in previous work (D’Amario et al., 2024; Tsay et al., 2024; Townsend et al., 2026), would be discovered. Although we illustrate three representative strategy time courses in Figure 5 (Aha! moment, incremental, and exploratory), we do not assume that all participants belong exclusively to three categories. Instead, these patterns were selected because they capture fundamental characteristics of strategy development. We suspect that a wide range of observed behavioral trajectories can be described through combinations or varying degrees of these underlying characteristics.

Eight standardized features (see table 1) capturing behaviour observed in the aiming trajectory were tested. Behavioural features such as baseline-normalized variability, learning duration, absolute trial-to-trial changes, and the proportion of direction flips in the aiming trajectory were used to index exploratory behavior, with higher values reflecting greater variability, reduced stabilization across trials, and frequent reversals in movement direction. Other features including learning duration and jump indices were incorporated which aimed to differentiate insight-like learning as short learning periods and an overall aiming trajectory comprising of one large abrupt change. Finally, incremental learning was characterized using measures of continuity in the aiming trajectory, including the coefficient of determination (R²) of the performance trend and trajectory smoothness. Higher R² values indicated more linear, improvement over time, while greater smoothness reflected incremental, continuous adjustments in aiming behavior. Table 1 presents equations for each feature, where *N* represents the total number of trials within the learning window (strategy development duration). Let *t* represent trial number, the trial-to-trial aiming change is defined as:

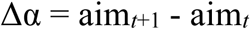

**Table 1.**
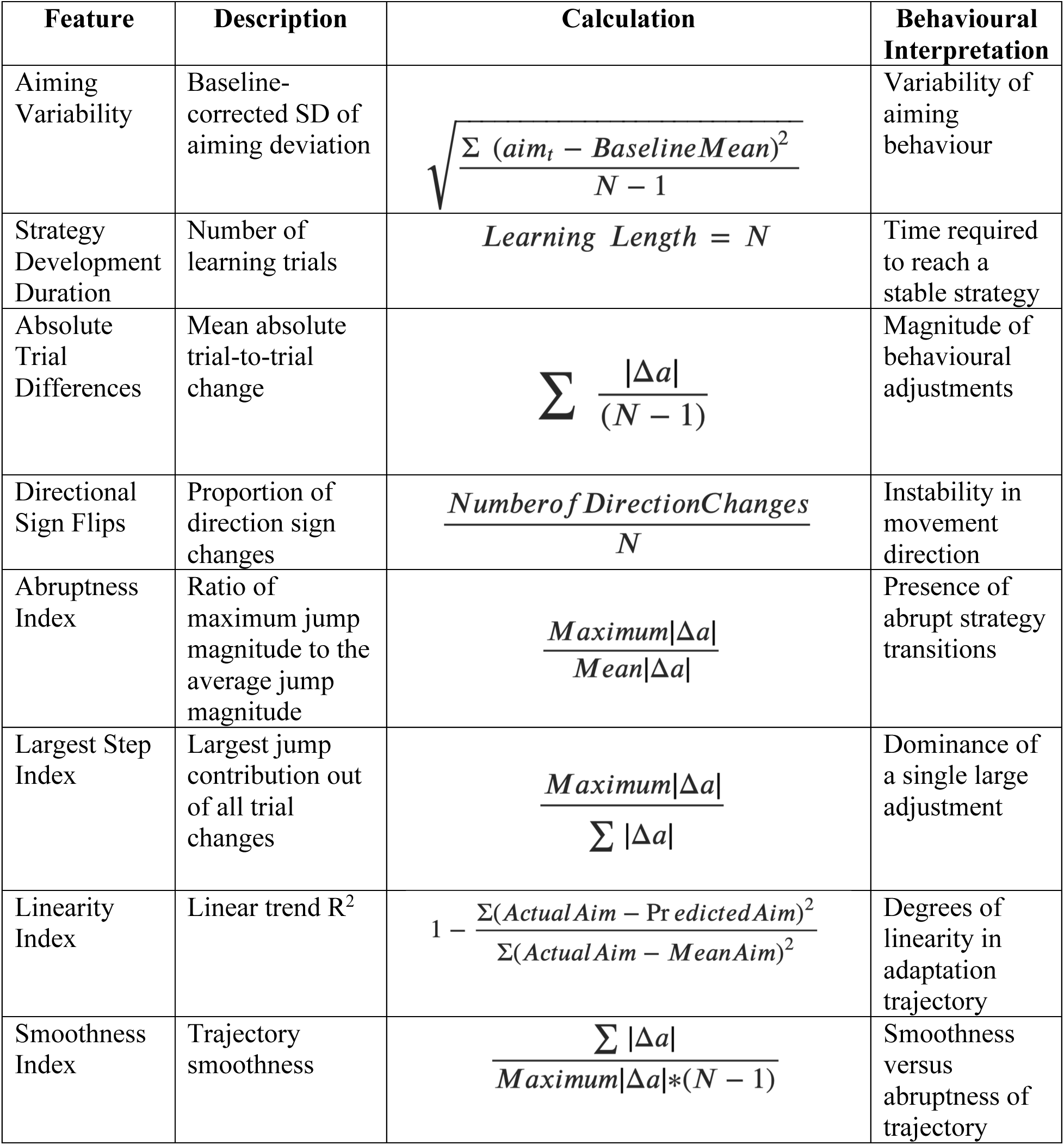
Summary of Extracted Learning Features. List of eight features for PCA input which describe observed behavioural patterns in the explicit timecourse.

A Principal Component Analysis (PCA) was run on these eight features. To determine which principal components were most informative, we used a scree plot to inspect the number of components that explained 80% of the variance (Joliffe & Cadima, 2016; Radeef et al., 2024). A linear regression will be used to assess if and how the most informative principal component scores vary with rotation size.

We next used a hierarchical clustering algorithm (HCA) to group types of strategy development timecourses. The HCA was performed on the most informative PCA components, where Euclidean distance was used as the dissimilarity measure. The number of clusters was decided by computation of a gap statistic (Zambelli, 2016), from visual inspection of the hierarchical dendrogram, and to align with the expected number of clusters (e.g. Tsay et al., 2024). Cluster labels will be determined by visually inspecting the explicit timecourses of participant representatives placed at the cluster centre. If representative participants display similar patterns in aim deviation, the assigned cluster label will be confirmed. Furthermore, generalized linear models (GLMs) will be implemented to further examine the relationship between rotation size and cluster separation. We will compare an alternative model containing the rotation predictor against an intercept-only null model using Akaike Information Criterion (AIC). While the null model assumed a uniform baseline probability of strategy use across all conditions, the alternative model allowed the likelihood of strategy adoption to vary as a function of rotation size.

Finally, implicit adaptation will be assessed using reach aftereffects during post-rotation no-cursor trials. Unlike the preceding analyses of explicit strategy development, analyses of reach aftereffects will include both strategy users and non-strategy users. If explicit and implicit adaptation are in competition with each other (Albert et al., 2022), greater explicit strategy use should be associated with smaller aftereffects. Conversely, participants who took longer to develop an explicit strategy may exhibit larger aftereffects, as implicit adaptation has more time develop before explicit compensation is fully established. First, a one-way ANOVA will first used to determine whether the magnitude of reach aftereffects differ based on rotation size. If a significant effect of rotation size exists, the aftereffects will be normalized by rotation magnitude. We will then test whether aftereffect magnitude differs between participants who did and did not develop an explicit strategy. Finally, among participants classified as strategy users, a one-way ANOVA will assess whether aftereffect magnitude differs across strategy- development groups.

## Results

### Adaptation and Aiming magnitude across rotation groups

Before investigating aiming timecourses, we first looked at overall adaptation (Fig 8a).

**Figure 8.**
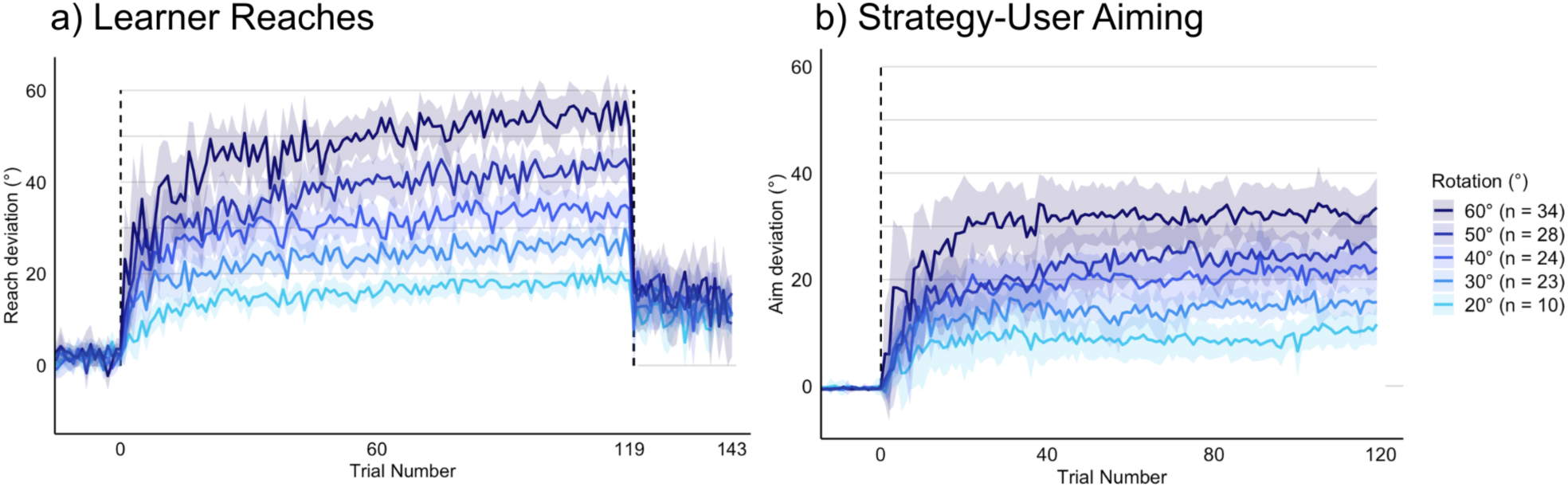
Average adaptation and strategy learning curves. **A:** Mean reach deviation (°) of learners across trials for the five rotation groups (20°, 30°, 40°, 50°, and 60°, N = 40 each group). The dashed vertical lines indicate perturbation onset and removal. Trials prior to this line reflect the aligned baseline performance. Shaded regions represent ± SEM across participants. **B:** Mean Aim deviation of strategy-users across trials for five rotation groups.

As expected, the magnitude of overall adaptation varied as a function of rotation size (*F*(4,195)= 127.4 *p* <0.001), with larger visuomotor rotations eliciting greater adaptation as shown in Figure 8. This result confirms that overall adaptation scaled appropriately with perturbation magnitude.

We then classified participants with and without strategy. Participants were classified as non-strategy users if the lower bound of the 95% confidence interval of their aiming responses during the last 16 perturbation trials did not exceed 5°, indicating that their aiming responses remained within the range of motor noise. We additionally excluded one participant who failed to stabilize, as indicated by sign flips relative to the target during the final 16 trials. Applying these criteria, 118 learners (59%) were classified as strategy-users (see Figure 8B, for the number of strategy users in each rotation condition). As with overall adaptation, we then looked at the group-average aiming time courses (see Fig 8B) and confirmed that the magnitude of final aim deviations varied with rotation size (*F*(4, 195) = 26.7, *p* < 0.001). To better visualize differences between overall reaching and aiming timecourses, we first plotted two-dimensional (2D) histograms containing all participant trial-by-trial responses. Whereas reach deviations formed the smooth, exponential-like pattern typically associated with visuomotor adaptation (Fig. 9A), aiming responses exhibited substantially greater heterogeneity, with participants displaying a wide range of temporal profiles, including abrupt transitions and more gradual changes (Fig. 9B; see also Fig. 4). We next sought to characterize the nature of this heterogeneity in explicit strategy development.

**Figure 9.**
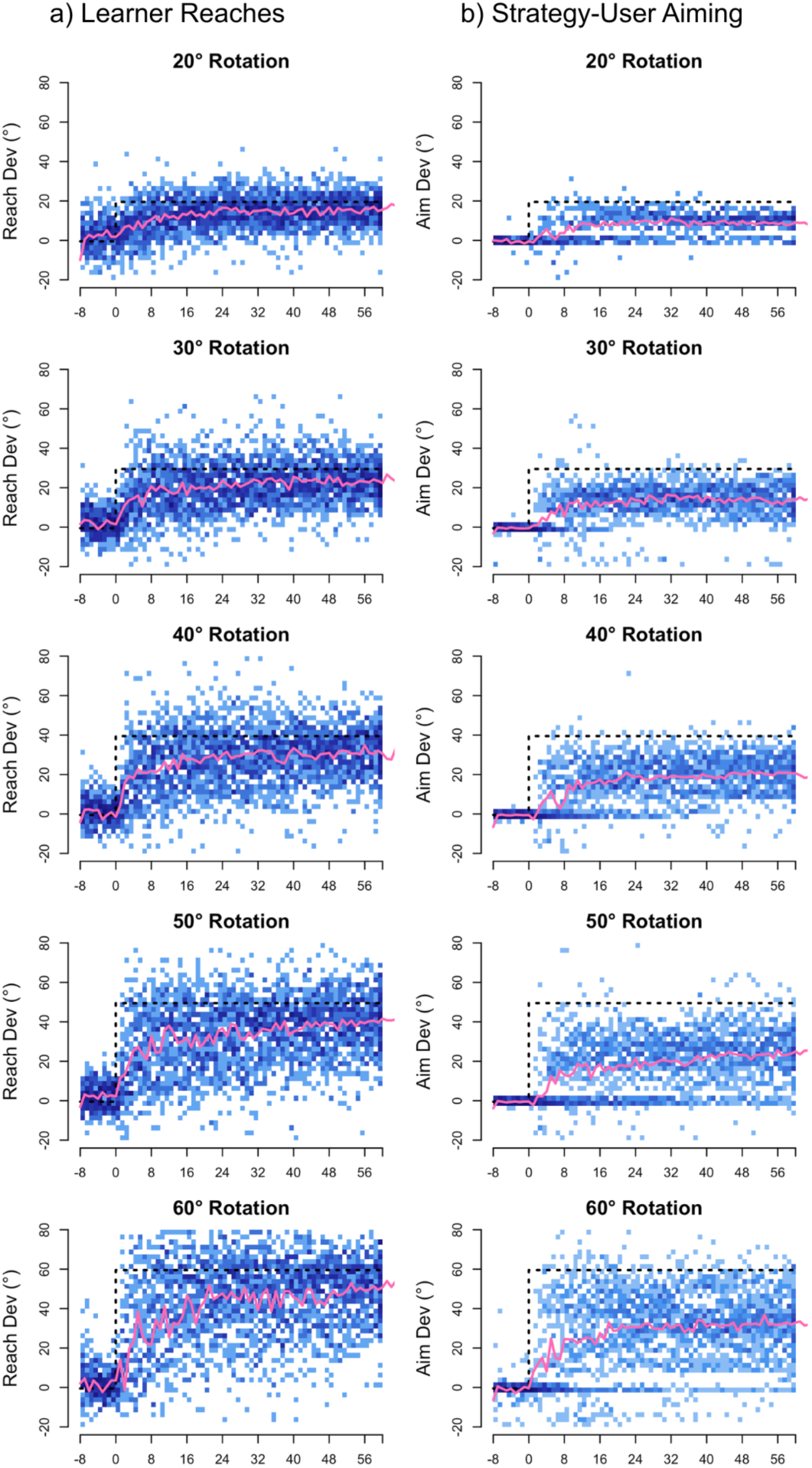
2D Histogram plots of trial-by-trial responses. **A:** Histograms containing individual reach deviations across trials for each rotation groups. **B:** Histograms of individual aim responses across trials in all rotation groups. In both panels, the dashed line indicate the perturbation onset (trial 1) and imposed rotation magnitude for each group.

## Defining Heterogeneity in Explicit learning

Previous data from our lab shows strategy development proceeding as a stepwise process (D’Amario et al., 2024). However, we observed different strategy development time courses in the current study that may not be explained by a stepwise function. Instead, some participants increased their strategy more incrementally or showed large trial-to-trial aiming changes in both directions for a longer period of time. Similar patterns have been found by others (Tsay et al., 2024). To incorporate this, we describe explicit strategy development, using a set of 8 features (see Table 1), and then applied dimensionality reduction with a Principal Component Analysis (PCA).

To characterize how behavioural features varied with rotation magnitude, Kruskal–Wallis tests were conducted across rotation groups, as shown in Figure 10. We note that aiming variability, strategy development duration, absolute trial differences, and largest step index features display statistically significant associations with rotation (all p<0.04). While for some of the features we might expect some effect of rotation size a priori, any variation associated with rotation may still reflect genuine differences in strategy development over time. Since characterizing behavioural heterogeneity was a primary focus of the present study, all eight features were retained for PCA. Figure 11A shows the cumulative variance explained by successive principal components. The first three components together, could explain about 79.96% of the total explained variance and were therefore retained for subsequent analysis.

**Figure 10.**
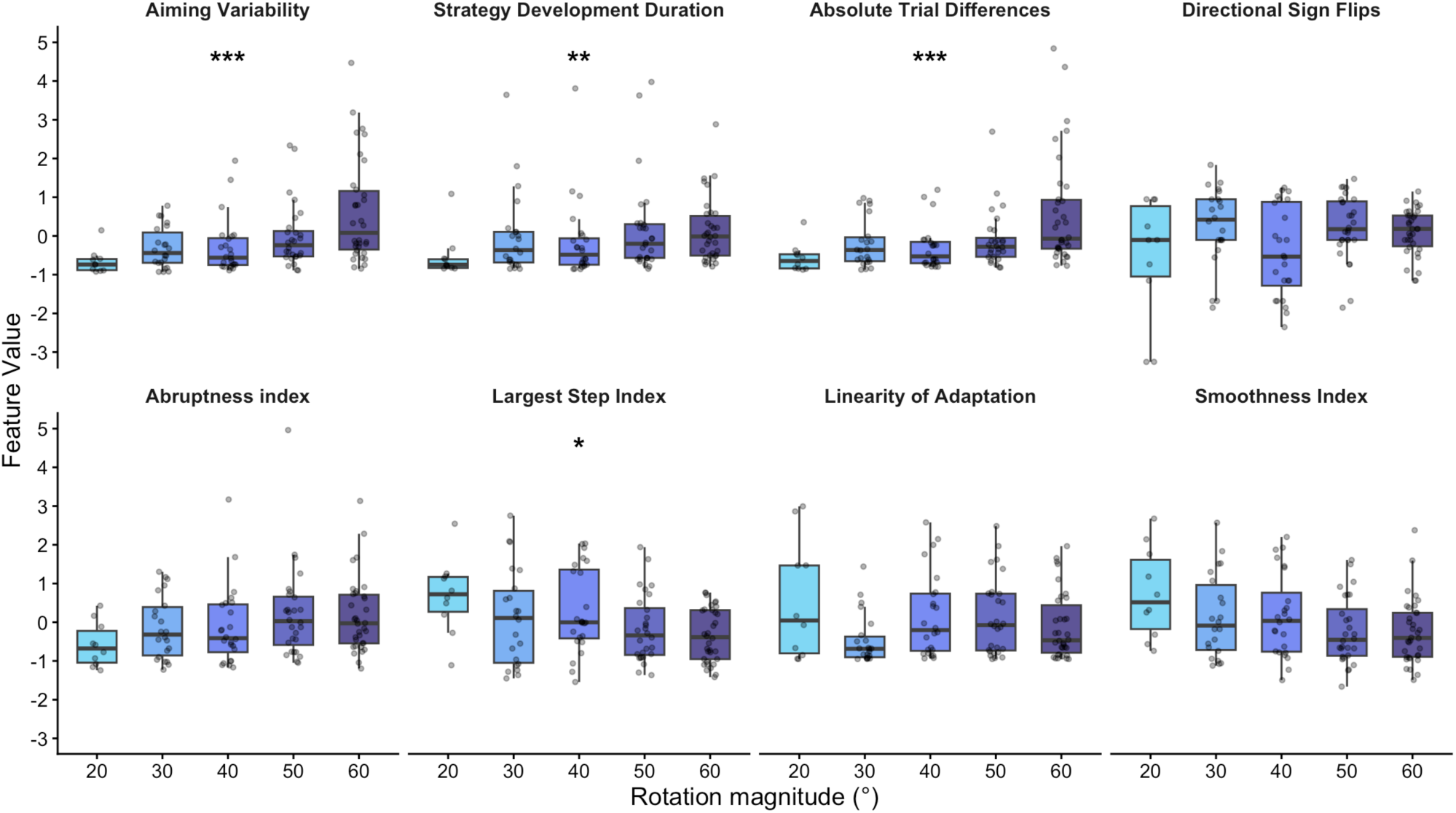
Feature Boxplots. Distribution of eight behavioral feature values across rotation groups, ordered from lowest (light blue) to highest (dark blue) rotation magnitude. Each dot represents an individual participant’s feature score. Asterisks at the top of a panel indicate a significant association (p<0.05) between rotation magnitude and that specific feature.

**Figure 11.**
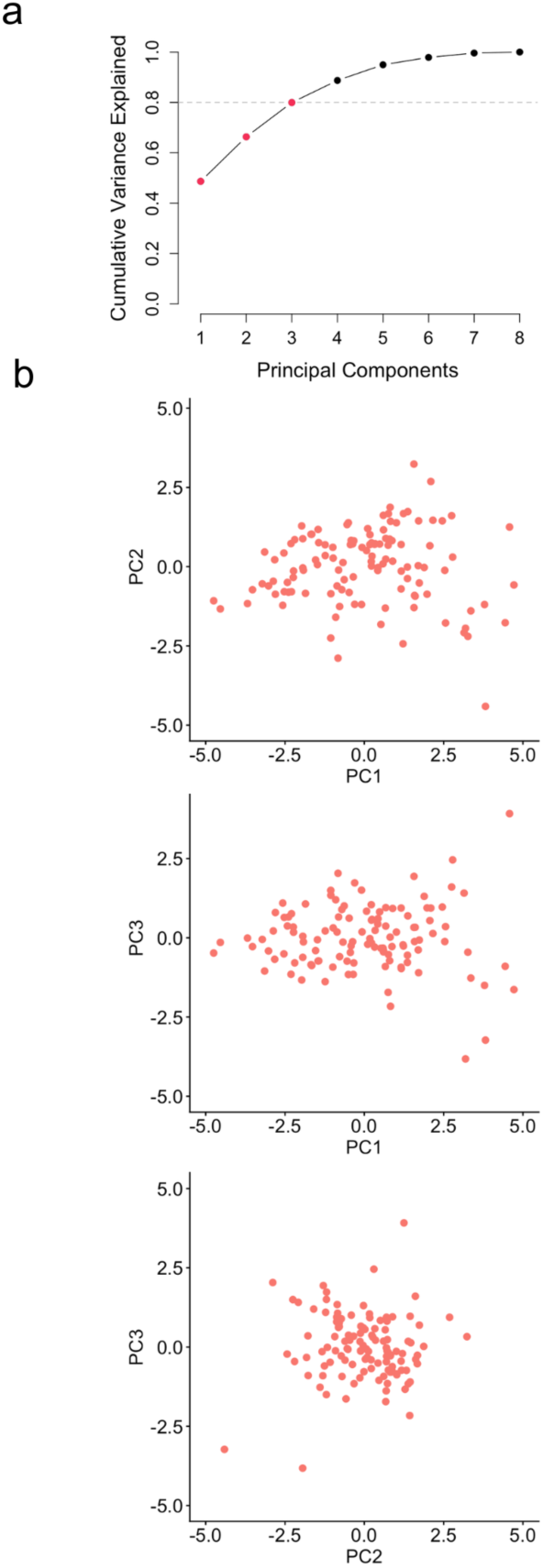
Most Informative Principal Components. **A:** Cumulative Variance Explained by All Principal Components. Dashed line represents 80% threshold. Red dots represent the principal components used for the hierarchical clustering analysis. **B:** Scatter plots showing the relationship between each pair of chosen principal components. Each dot represents a participant projected into the PCA space.

Inspection of the component loadings (Table 2) indicated that PC1 primarily reflects a smooth- versus-variable learning dimension with higher scores associated with greater with aiming variability, and larger trial-to-trial adjustments. In contrast, lower PC1 scores are associated with smoother, more linear aiming trajectories. PC 2 and 3 both appeared to capture aspects of adaptation rate. PC2 contrasted trajectories characterized by large, single-step adjustments with those in which aim changes were more distributed over time. PC3 differentiated smoother, incremental adaptation from adaptation driven by discrete jumps.

**Table 2.** PCA Feature loadings.

| Feature | PC1 | PC2 | PC3 |
| --- | --- | --- | --- |
| Aiming Variability | 0.37 | -0.50 | -0.30 |
| Strategy Development Duration | 0.36 | 0.33 | 0.16 |
| Absolute Trial Differences | 0.36 | -0.40 | -0.49 |
| Directional Sign Flips | 0.35 | 0.31 | -0.05 |
| Abruptness Index | 0.37 | -0.20 | 0.56 |
| Largest Step Index | -0.33 | -0.46 | 0.30 |
| Linearity Index | -0.30 | -0.31 | 0.16 |
| Smoothness Index | -0.40 | 0.18 | -0.46 |

Figure 11B illustrates participants’ distributions within the three-component PCA space using pairwise projections. While distinct clusters are not visually apparent, PC1 appears to capture the largest spread in samples. The distribution along PC3 shows overlap, and no additional cluster structure is apparent in the PC1–PC3 or PC2–PC3 scatterplots. Therefore, the following analyses will be primarily focused on the PC1–PC2 plane.

## Aiming behaviour as a continuum

Although PCA space did not reveal distinct, well-separated clusters (Figure 11B), participants did not appear to be randomly distributed throughout the PCA. Instead, explicit strategy development appeared to vary along a continuum, with individual timecourses occupying intermediate positions that combined characteristics of the hypothesized strategy profiles. When we look at the mean and standard deviation along PC1 and PC2 (Fig 12A) this interpretation is further supported. We can see a gradual shift along PC1 in the group centroids across rotation conditions, although there is substantial overlap between neighboring rotation groups (Fig. 12A). To determine whether this continuum was systematically related to perturbation size, we next examined whether rotation size predicted participants’ PC values using a linear regression. These analyses indicated that rotation size significantly predicted participant scores on the first principal component (*F*(3) =11.34, *p* =.044, *r* =.33 see Fig 12B), but not PC2 (*F*(3) = 1.21, *p* = .35, *r* = -.17) or PC3 (*F*(3) = 0.41, *p* = .57, *r* = -.11). Together, the progressive shift in group centroids (Fig 12A) and the significant relationship between rotation size and PC1 (Fig 12B) support a graded organization of explicit strategy development, whereby increasing perturbation magnitude shifts participants progressively along a continuum of strategy-development profiles.

**Figure 12.**
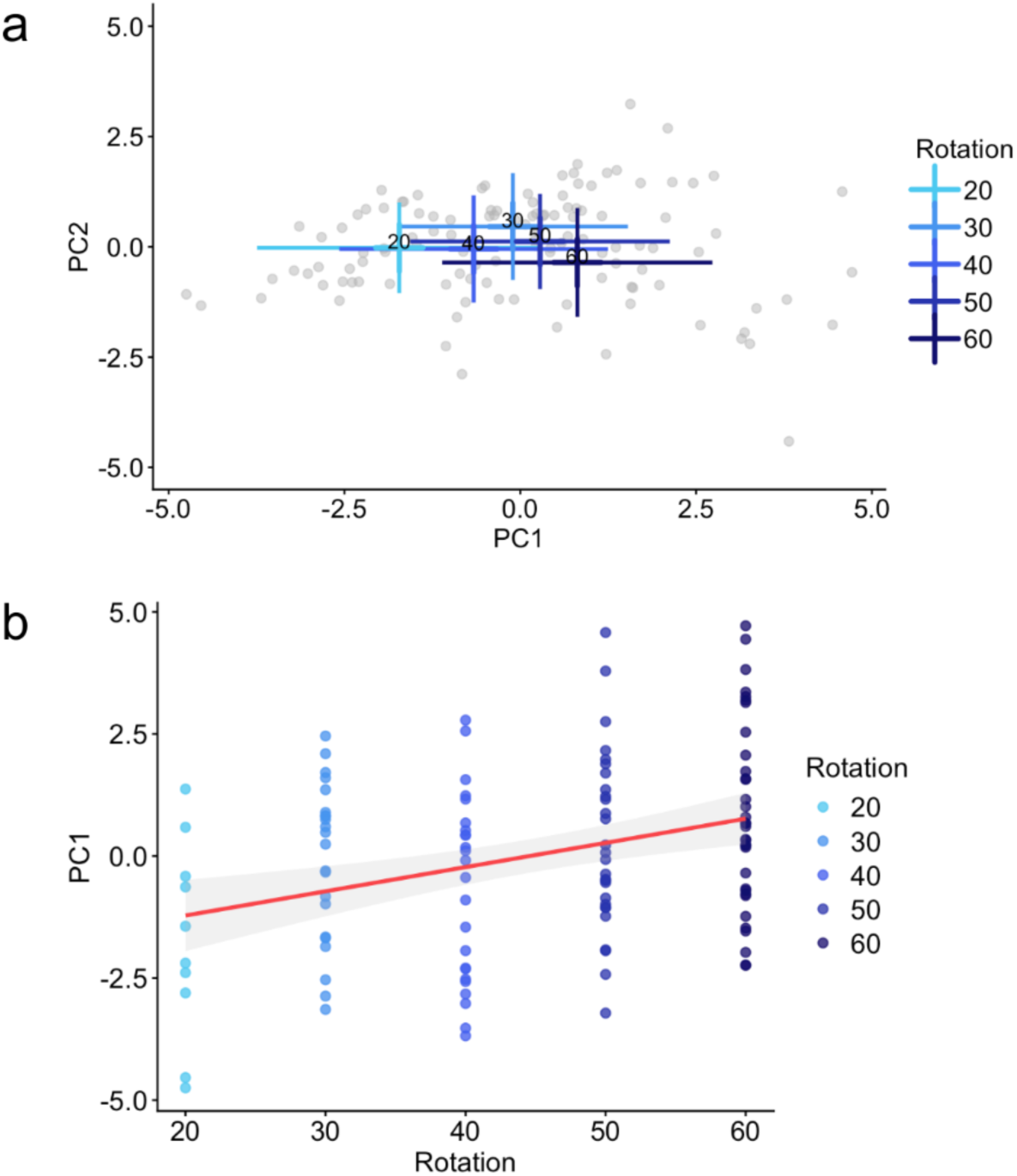
Explicit Learning as a Function of Rotation. **A:** PC values per rotation group across the PCA space. The centroid of each plus sign represents the group’s mean, and the orthogonal line extensions represent the standard deviation. Grey dots represent participants within the space. **B:** PC1 and Rotation Linear trend. Each point represents a single participant’s PC1 score. The red line represents the fitted linear regression line.

## Explicit strategy-development clusters

Despite the absence of visually distinct clusters in the PCA space, previous work has proposed discrete strategy-development types (D’Amario et al., 2024; Tsay et al., 2024; Townsend et al., 2026). We therefore applied a hierarchical clustering algorithm (HCA) to determine whether similar groupings emerged in our data. The first three principal components were used as input to the HCA. The gap statistic reached its maximum value at *k* = 3, and inspection of the dendrogram indicated that a three-cluster solution was appropriate (Fig. 13A).

**Figure 13.**
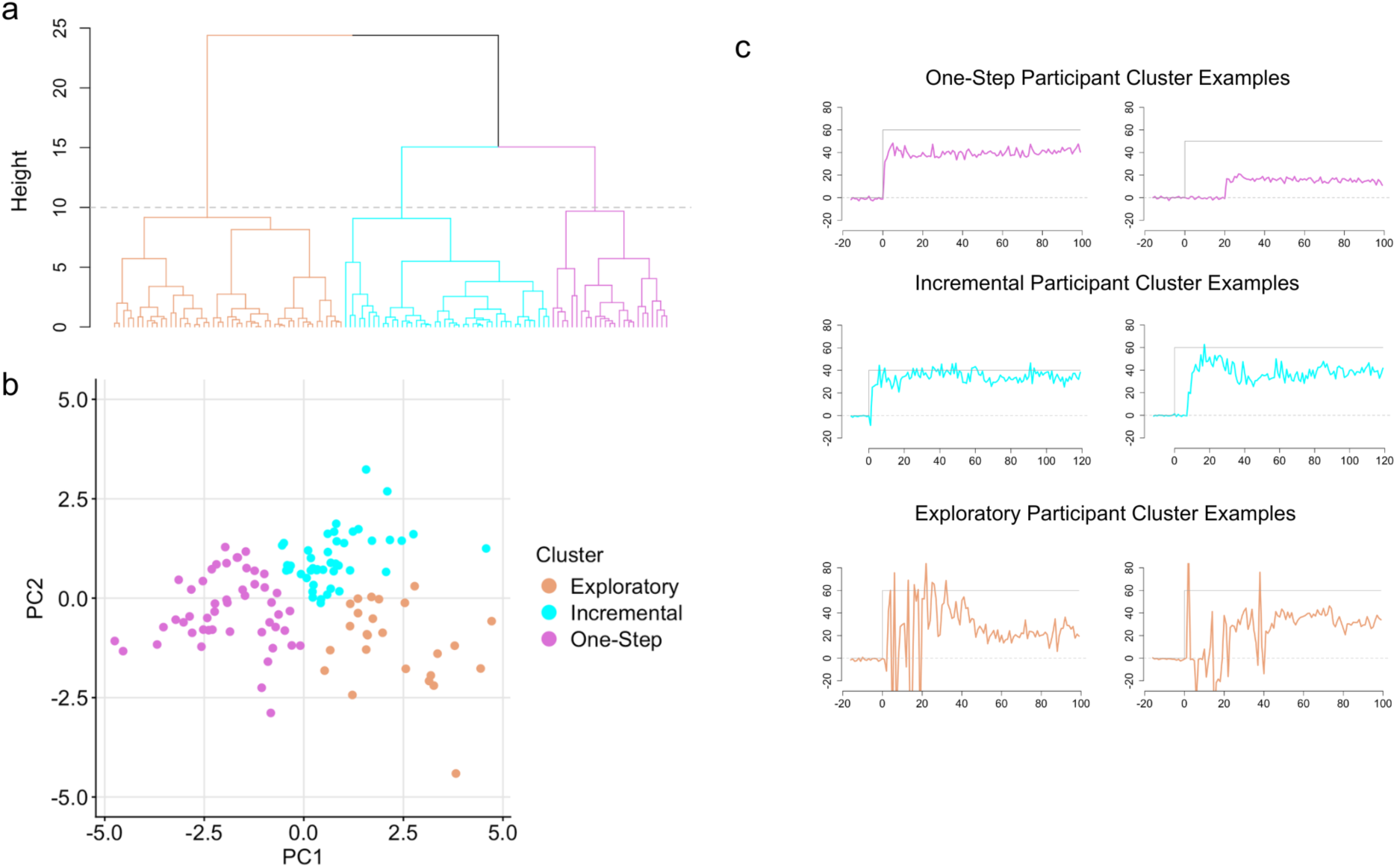
Hierarchical Clustering Results. **A:** Hierarchical Dendrogram Structure. Dashed line represents height cutoff. Each colour represents a cluster group. **B:** Clusters across the PCA space. PC1 represents trajectory volatility with smoother, more linear trajectories at the lower end. PC2 distinguishes the nature of the variance, with lower values capturing large steps and higher values reflecting distributed, incremental adjustments. **C:** Example participant aiming trajectories derived from each cluster centroid.

The resulting clusters of strategy development occupied neighbouring regions of PCA space (Figure 13B). Clusters were labeled based on visual interpretation. One cluster was characterized by exploratory behaviours (n = 25, orange in Figure 13), in which participants sample a wide range of aiming solutions in various directions before settling on a strategy. Next, we find some strategy-users can almost immediately settle on an aiming strategy, which we label as stepwise learning (n = 49; pink in Figure 13). Finally, others instead implement a more graded approach sometimes showing multiple abrupt changes in aim early in learning. We therefore refer to the latter cluster as incremental (n = 44, cyan in Figure 13).

Importantly, these clusters should not be interpreted as distinct categories of explicit strategy development. Rather, consistent with the continuous organization observed in the PCA space (Figs. 11B and 12), the clusters appear to represent neighboring regions along a broader continuum of strategy-development profiles.

We assessed whether differences in strategy cluster distribution changed across rotation conditions. Figure 14 displays these findings, visualizing the proportion of strategy-users and cluster distributions across rotation groups. Specifically, the ’Exploratory’ cluster (represented in orange) is absent at 20° (n = 0) and nearly absent at 30° (n = 2) but expands to become the dominant type at 60° (n = 13). We also see non-strategy-users (represented in white) shrink as rotation size grows. This progression visually underscores how larger explicit demands drive a transition from stable responding into more active behavioral exploration. To see if these group differences were meaningful, we fitted a multinomial generalized linear model (GLM) to determine if strategy development depends on rotation. Model support was evaluated using Akaike’s Information Criterion (AIC) by comparing the fitted model to a null model containing no rotation effect. Evidence for the alternative model was inferred when the difference in AIC between models (ΔAIC) exceeded 2 (e.g. Burnham & Anderson, 2004). A binomial GLM was used to model the proportion of participants utilizing a strategy versus a non-strategy. Comparing a model with a rotation effect against the null model with one proportion, yielded a ΔAIC of 29.6, providing evidence that likelihood of strategy development varied as a function of rotation. Next, we fitted a multinomial GLM, offering insight into the critical role of rotation size on cluster type. The alternative model, which included rotation size as a predictor, significantly outperformed the null intercept-only model (ΔAIC=7.7). Additionally, we computed binomial GLMs to each strategy cluster as a post-hoc measure, revealing that exploratory (ΔAIC = 9.4), but not single and incremental phenotypes depend on rotation size. While the clusters identified here, do not seem to correspond to discrete classes of strategy development we do observe a strategy continuum in which participants develop an aiming strategy. Overall, the way participants develop their strategy varies as a function of rotation size. Specifically, with larger rotations, participants seem to explore the solution space more, corresponding to both a higher value on PC1 and hence higher prevalence of the “exploratory” cluster.

**Figure 14.**
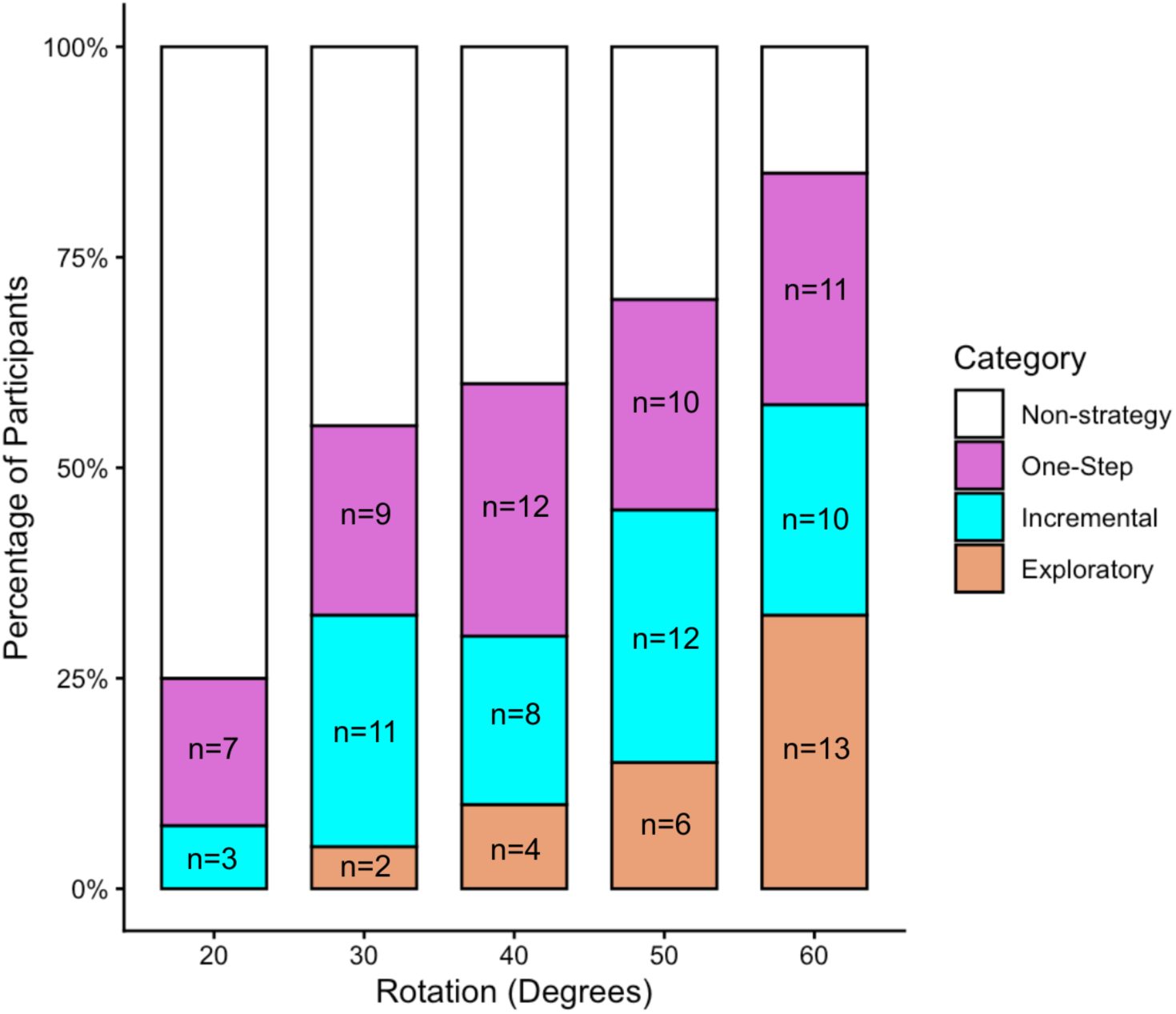
Distribution of Strategy Development Phenotypes Across Rotation Groups. Stacked bar plot displays percentage of participants classified into distinct strategy groups (Non-strategy, One-step, Incremental, Exploratory) as a function of rotation size. Sample sizes for each sub- group are overlaid each bar. As rotation increases, exploratory strategies become more prevalent while non-strategy behaviours decrease.

## Effect of explicit learning heterogeneity on implicit aftereffects

Following the rotation phase, participants completed no-cursor reaches feedback to assess implicit aftereffects. Figure 15 illustrates the average no-cursor reach deviations, separated by rotation (A), strategy use (B), and strategy development cluster (C). Because rotation size determines final reach adaptation in the rotation phase (See figure 8), we first set out to determine if reach aftereffects (based on the average of the first 8 trials) differ by rotation. These reach aftereffects were significant (deviations were above 0 (*t*(199) = 9.7, p < .001) but did not vary with rotation size (*F*(4, 195) = 0.58, *p* = .68) nor by use of strategy or not (*t*(122.8) = 1.05, *p* = .32). We set out to determine if implicit aftereffects differed across strategy development clusters (for those participants showing a strategy). Although participants assigned to the exploratory cluster tended to exhibit slightly larger aftereffects than participants assigned to the incremental cluster (Fig. 15C), these differences were not statistically significant (*F*(2, 115) = 0.90, *p* = .41). Overall, implicit aftereffects were observed across participants, but were not significantly influenced by rotation size, explicit strategy engagement, and assignment to a strategy development cluster. This means that in this data set, implicit reach aftereffects seem to be independent of explicit strategy.

**Figure 15.**
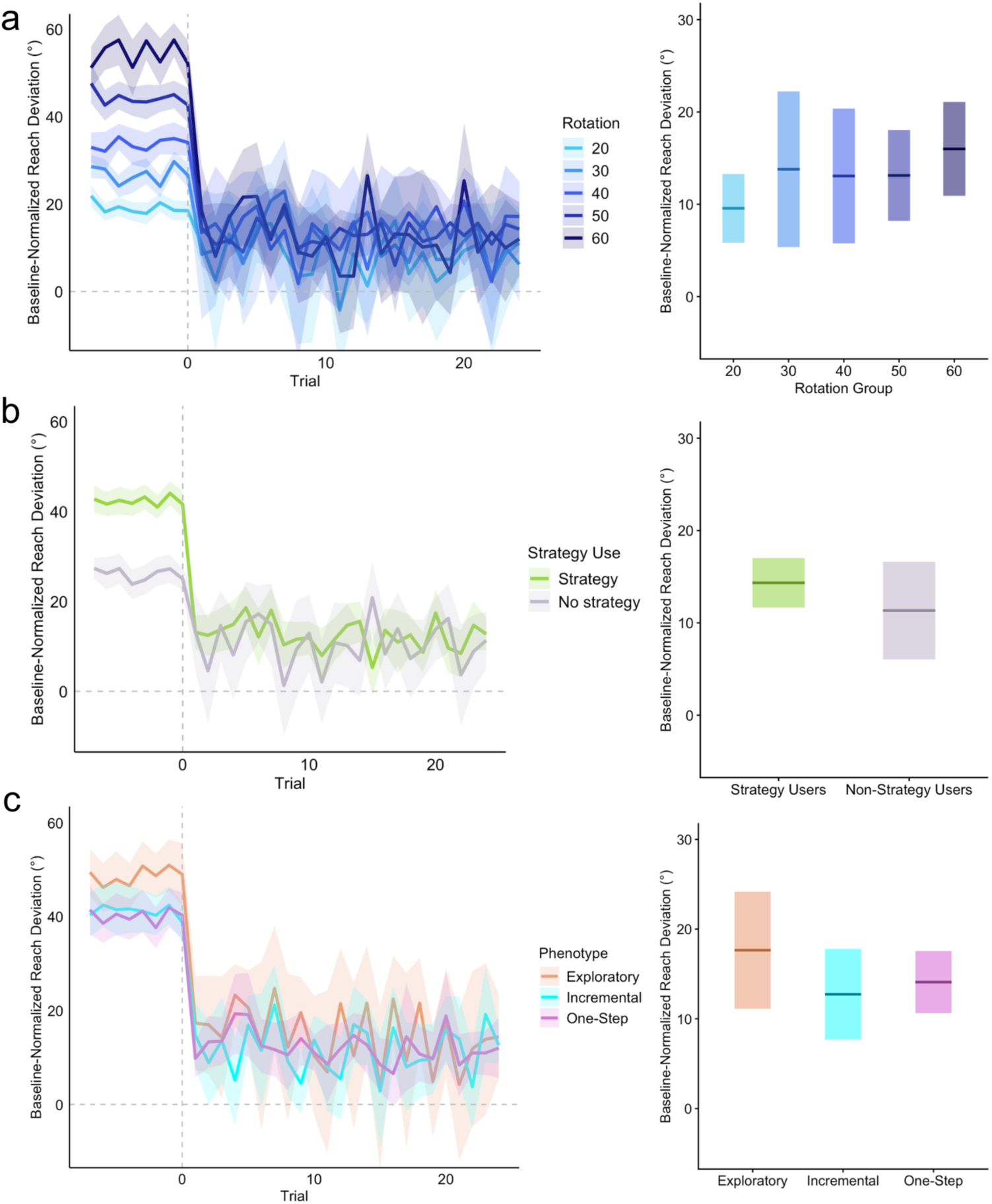
Reach aftereffects. **A:** Average no-cursor reach across rotation groups shown in the left panel. The right panel shows 95% confidence interval boxplots. Darker horizontal lines reflect group mean, **B:** Average no-cursor reach across strategy and non-strategy learners.

Dashed vertical line represents final trial before cursor feedback removal. Shaded regions represent ±SEM across participants. **C:** Average no-cursor reach across strategy clusters.

## Discussion

Here, we demonstrate that explicit strategy development is not a homogenous process, nor can it be neatly compartmentalized into a few subtypes of strategy development timecourses.

Instead, we found that explicit strategy development lies along a continuum, spanning from sudden stepwise moments of insight to prolonged exploration of the solution space, as well as many intermediate forms. We further find that perturbation size influences strategy learning. First, the proportion of participants learning a strategy increases with rotation sizes. Second, the mix of strategy learning timecourses changes with rotation size. Participants exposed to larger perturbations learn in more exploratory patterns.

A recent study (D’Amario et al., 2024) has suggested that explicit strategies often emerge through a discrete moment of insight, producing abrupt transitions in aiming behaviour that resemble a stepwise pattern. Their study (see Fig. 4 of present study and Fig. 10 in D’Amario et al., 2024) and work from Townsend et al. (2026) compared stepwise and continuous learning-curve models, and found that stepwise changes provided a better description of participant aiming performance. Our findings are somewhat consistent with these observations. Indeed, one prominent region of the strategy-development continuum was characterized by abrupt transitions into stable aiming solutions. However, our results further suggest that stepwise strategy acquisition represents only one region of a broader space of explicit learning behaviours. We observed additional regions of the continuum characterized by a more volatile or incremental strategy development where aiming progressed through a series of abrupt transitions before stabilizing, as well as exploratory patterns in which multiple possible solutions were tested before converging on a strategy. Many participants were found to exhibit intermediate forms of the three clusters, blending characteristics of two or more cluster types. The clusters we identified should probably be viewed as regions along the continuum, rather than discrete, mutually exclusive categories.

This broad interpretation is consistent with recent findings (Tsay et al., 2024), who highlighted substantial variability in explicit aiming behaviour across visuomotor adaptation studies, including fast and slow insight, systematic and erratic exploration. The behavioural patterns described by Tsay et al. (2024) resemble the regions of the strategy continuum identified in the present study. Rather than supporting a single dominant mode of explicit learning, our findings suggest that explicit strategy development may span a continuous range of behaviours that differ in how strategies are discovered, evaluated, and refined over time.

Although the present results indicate that explicit strategy development varies continuously across individuals, they do not explain *why* participants adopt different learning trajectories. One possible explanation is that individuals differ not only in how quickly they discover a strategy, but also in how they represent the visuomotor perturbation itself. This possibility is consistent with the framework proposed by Ding et al. (2026). Rather than only studying the temporal development of explicit strategies, Ding and colleagues used the full adaptation timecourse and also included the content of participants’ learned strategies by assessing how strategic rules generalized to novel probe-target locations during and following adaptation. Their findings suggest that individuals may differ in how they represent the relationship between actions and outcomes, resulting in different strategic rules even when behaviour at the trained targets is similar. Since participants reached to a probe every third trial, probe-target measures do not provide as direct a measure of explicit strategy development as continuous aiming reports. These findings raise the possibility that some of the heterogeneity observed in the present study reflects differences in how participants understand and counter the perturbation.

In contrast, the aiming reports in our study allow the development of explicit learning to be quantified on a trial-by-trial basis, providing a more direct measure of the strategies individuals choose and how they evolve throughout the timecourse. While we do not probe the content of the strategy, our approach offers great insight into the temporal dynamics of strategy development. Although our study cannot directly distinguish between different internal representations, the hypothesis-testing framework proposed by Ding et al. (2026) provides a useful lens through which to interpret the behavioural diversity observed here and Tsay et al., (2024).

Within this framework, participants exhibiting abrupt transitions may have rapidly inferred a successful compensatory rule, producing the stepwise strategy-development patterns described by D’Amario et al. (2024) and Townsend et al. (2026). Elsewhere along the continuum, participants exhibiting exploratory may have evaluated completing candidate solutions before converging on a stable strategy. Intermediate, incremental patterns may reflect successive adjustments of an initially incomplete solution, producing the series of abrupt transitions observed in the incremental region of the strategy continuum. Importantly, these possibilities are not mutually exclusive. Rather, different regions of the strategy continuum may reflect varying combinations of insight, exploration, hypothesis testing, and strategic refinement.

Taken together, these findings suggest that a single account of explicit strategy development is insufficient to explain the diversity of strategy-development behaviours observed during visuomotor adaptation. Rather than representing a discrete set of strategy types, explicit strategy development appears to lie along a continuum spanning abrupt insight, iterative refinement, and extended exploration. Importantly, perturbation size systematically shifted where participants fell along this continuum, with larger perturbations increasing both the likelihood of explicit strategy use and the prevalence of more exploratory forms of strategy development. Compared to implicit and overall adaptation, explicit strategy development appears substantially more heterogeneous, with individuals differing not only in whether they develop a strategy, but also in how that strategy emerges and stabilizes over time.

## Rotation size as a mechanism for modulating explicit heterogeneity

How might strategy development timecourses therefore vary across participants? Our data shows two effects of rotation size. First, the number of people learning a strategy increases with rotation size. Larger perturbations may increase error saliency and this may guide a larger proportion of participants to develop a strategy (Modchalingam et al., 2019; D’Amario et al., 2024). Our findings confirm that perturbation salience may increase the probability that each participant develops an aiming strategy.

Second, rotation size was associated with changes in trajectory smoothness, variability and abruptness level, aligning with shifts in participant positions along the first principal component. While the data indicate a continuous distribution, primarily dictated by the first principal component, we can nevertheless use the clusters to understand the continuum. The changes in cluster prevalence over rotation size (Fig 14) are a direct consequence of the shift along the first principal component (Fig 12). Since the proportion of participants who develop a strategy also increases with rotation size, this means that participants who did not develop a strategy in the smaller rotations, may have been driven into developing a strategy with an exploratory timecourse had they been presented with a larger rotation. In other words, the saliency of errors may drive if and how participants develop an explicit strategy.

## Cognitive capabilities as a mechanism for explaining explicit heterogeneity

Stepwise and incremental patterns of strategy development were present across all rotation magnitudes, suggesting that these forms of explicit learning may emerge relatively independently of perturbation size. This raises the question why many participants identify strategies by engaging in very little exploration. One possibility is individual differences in working memory, rather than task demands alone. For example, motor working memory capacity has recently been shown to positively predict explicit adaptation (Hillman et al., 2025).

Finding a strategy solution in one or, a few steps, may be beneficial, reducing the need for unnecessary exploration and noise in performance. Christou et al. (2016) highlight a subset of participants with verbal aiming reports in the wrong direction, observing that these individuals had low working memory scores. According to these authors, individuals who possess high working memory capacity are equipped with maintaining and evaluating multiple solutions simultaneously. In the present study, participants are exposed to an 8-target workspace, likely exceeding cognitive capacity load if a separate solution/response would be developed for each target (McDougle & Taylor, 2019). This number of targets was chosen to drive global (“parametric”) strategy development in most – if not all – participants. Nevertheless, a high working memory capacity may facilitate retention from previous trials. This can allow individuals to remember prior aiming outcomes and quickly evaluate current performance for deliberate adjustment. This ability may be supported by a greater efficiency in integrating prior information and guide spatial transformations, as previous work has linked early visuomotor learning rate with mental rotation performance (Anguera et al., 2010). In contrast, individuals with lower working memory capacity may face challenges in maintaining previous trial information, leading to greater reliance on trial-and-error exploration. Differences in working memory capabilities are also noted across both younger and older adults, with higher working memory being associated with greater explicit strategy development (Vandevoorde & Orban de Xivry, 2020). Directly testing how working memory capacity influences where individuals are positioned in the strategy-continuum remains an important direction for future work.

## Other potential mechanisms

Furthermore, these differences in cognitive resource recruitment may be shaped by measures of task success (i.e., incentive to develop a well-defined strategy). Some participants may value precision in their aiming solution, requiring greater cognitive engagement rather than a rough approximation of the solution space. A recent study by Yamada et al (2025) offers an idea that individuals who hold cognitive biases that allow them to be less confident in their decision making, show more exploration in their explicit strategy, such that trial-by-trial changes and variability in re-aiming are larger. These authors also suggest that engaging in a larger re-aim strategy might allow participants to receive enhanced clarity in feedback. Therefore, non-exploratory strategies, may allow participants rapidly adopt a global rule (e.g., “aim counterclockwise”), whereas those engaging in more exploratory strategies might be reflecting ongoing investigation of the rule space to identify the most optimal strategy.

## The importance of task understanding

We find people who consistently aim at the target throughout the task, even in the 60- degree group. This may be because those participants still did not understand what they were supposed to do during the arrow aiming task. Task understanding is found to be needed for proper explicit/implicit component isolation. Chen & Taylor (2025), specifically found weaker implicit-explicit associations of individuals who did not understand task instruction, such that they consistently aimed at the target during strategy inclusion trials. Other work has noted the need to provide participants with additional instructions or guidance during re-aiming (Wilterson & Taylor, 2021), proving it difficult to probe the spontaneous timecourse of explicit learning.

While self-aiming the arrow offers insight into participants’ planning throughout the task, the recorded adjustments may not perfectly reflect the cognitive process. We decided not to provide any additional instructions after the rotated phase had started, to really be able to record the natural time course of strategy development. For some fraction of participants, this may have led them to not report their actual hand-movement strategy, but instead report the intended cursor-movement - despite our extensive prior instructions as well as the demonstration of the difference between the hand and cursor using error-clamped trials. We did find non-strategy learners in all rotation groups, suggesting that some participants may have had an aiming strategy but did not express it when orienting the arrow. Anecdotally, some participants did mention strategies during debriefing, despite not reporting them with the arrow. Future work can implement post-experiment qualitative measures or cognitive tests to assess participant task-related planning and understanding of instructions to better determine if they did or did not have a strategy.

## Future Directions

The labeling of clusters was defined using representative participants at the centroid of each cluster (e.g. see Fig 13C). As a result, not all participants perfectly align with their predefined behavioural labels. These differences may reflect the idea that participant strategies exist along a continuum rather than in distinct categories. Participants might also be switching between multiple strategy phenotypes throughout the task. Nevertheless, the explanatory power of the features and PCA suggests that participants use various methods to develop or discover a strategy, which is influenced by perturbation size. As this work is one of the first to record, analyze and define individual strategy development timecourses, future work may be able to distinguish more informed cluster boundaries or switching between methods of strategy development.

Second, while we included a no-cursor reach block following adaptation, our paradigm was not designed to characterize the full timecourse of implicit adaptation. Instead, the focus was on identifying patterns in the emergence of explicit strategies. Consequently, we obtained only final reach aftereffects. Previous work suggests that implicit adaptation can develop relatively rapidly (Ruttle et al., 2021; D’Amario et al., 2024) and reaches a similar cap across a range of perturbation magnitudes (Bond & Taylor, 2015; Morehead et al., 2017; Modchalingam et al., 2019). Here we do not see any effect of the explicit timecourses on the final level of implicit adaptation. Nevertheless, interactions between implicit and explicit processes throughout learning may contribute to the observed behavioural patterns (Albert et al., 2022; ’t Hart et al 2024; D’Amario et al., 2024). Future work should determine any effect of the strategy development timecourse on the trial-by-trial timecourse of implicit adaptation.

Finally, while this study is one of the first to attempt to identify distinct strategy development styles, alternative approaches may provide complementary insights. Dynamic time-warping (DTW) was used in previous work (Ding et al., 2026; Tsay et al., 2024) to calculate dissimilarities between participants. Behavioural clusters were then showed to be separate in a t-SNE. Dynamic time-warping uses the entire explicit trajectory that is not reliant on behavioural features. In the present study, we instead extract behavioural properties of the aiming timecourse during the strategy development phase only, to identify discrete styles of explicit strategy development. Because dynamic time-warping incorporates information from the entire adaptation trajectory, some of the group differences identified in prior work may reflect behavior occurring before strategy development onset or after strategy stabilization. Future work could investigate the role of features that capture these phases of learning, such as strategy stabilization time, variability during stable performance, or normalized strategy magnitude. Using such features could align our approach with previous work, and perhaps this would show that previous work captures the properties of the strategy more so than the style by which the strategy was developed. Additionally, other feature sets might be able to identify distinct strategy development styles. However, if strategy development is not governed by separable cognitive styles, but indeed lies on a continuum, as our results suggest, this would be expected to produce a distribution whose properties vary as a function of rotation size. A descriptive model of the distribution of aiming responses depending on rotation size could in turn lead to a generative model and a deeper understanding of strategy development in motor adaptation.

## Conclusions

During a visuomotor adaptation task, implicit and explicit processes both reduce the discrepancy between expected and actual movement outcome following perturbation exposure. While implicit adaptation has been shown to be a homogenous, and gradual process, previous research has assumed the same for explicit processes as well. However, the current work shows that explicit strategy development is rather heterogeneous. Explicit strategy development ranges from stable or rigid to highly volatile or erratic, reflecting behavioural hallmarks of stepwise and exploratory patterns. This gradient is reflected in rotation size, such that larger rotations encourage more individuals to explore for alternative solutions, thereby increasing the likelihood that explicit strategies emerge. This suggests that the saliency of a perturbation affects both if and how strategies to deal with a visuomotor-rotation are developed. One consequence of these findings is that averaged explicit timecourses hide the heterogeneity in individual participants’ explicit strategy development styles, and hence the true nature of the explicit strategy development (D’Amario et al., 2024; Tsay et al., 2024). These findings inform the adoption of new methodologies and models that more accurately characterize the explicit component of visuomotor adaptation. This will contribute to the understanding of visuomotor adaptation as a set of processes.

